# Decoding Brain Interstitial Transport In Vivo: A Fully Validated Bottom-Up Mechanistic Prediction Framework

**DOI:** 10.1101/2025.03.22.643151

**Authors:** Tian Yuan, Nicolò Pecco, Wenbo Zhan, Stefano Galvan, Riccardo Secoli, Marco Riva, Lorenzo Bello, Andrea Falini, Ferdinando Rodriguez y Baena, Antonella Castellano, Daniele Dini

## Abstract

The transport of fluids and substances within the brain parenchyma (*i*.*e*. interstitial transport) is fundamental to maintaining brain health and delivering treatments for neurological disorders. However, accurately predicting these transport processes has remained a formidable challenge due to the intricate and dynamic nature of the brain’s microenvironment. Here, we report a novel, fully validated bottom-up mechanistic framework that bridges advanced mathematical modelling, ultra-high-resolution imaging, and biomechanical testing to achieve precise, *in vivo* predictions of interstitial transport. Using this approach, we accurately modelled the transport of MRI tracers in living sheep brains, offering unprecedented insights into the interplay between fluid dynamics and tissue properties. This platform is a transformative step forward, with the potential to revolutionise drug delivery strategies not only in the brain but also in cancer therapy and other soft biological systems. By addressing limitations in modelling complex transport in soft tissue, our work establishes a significant tool with profound implications for biomedical engineering and translational medicine.

---

The brain, often widely regarded as the most complex organ in the known universe, continues to present profound challenges to scientific inquiry Despite extensive research efforts, fundamental aspects of brain function remain elusive, including the mechanisms governing the transport of fluids and substances within the brain parenchyma (*i*.*e*. interstitial transport). This critical process has intrigued scientists for centuries [1–3] and gained renewed attention in 2012 with the discovery of the glymphatic hypothesis [4]. However, the precise mechanisms driving interstitial transport remain enigmatic, representing a significant knowledge gap in neuroscience.

Cerebrospinal fluid (CSF) and interstitial fluid (ISF) together constitute the fluid system of the brain [5]. CSF is primarily produced in the ventricle and stored in the subarachnoid space where it bathes the brain and spinal cord to protect them from mechanical shock and also allow them to maintain their functional shape [6]. According to the glymphatic hypothesis [4], CSF flows into the brain parenchyma via periarterial space and mixes with the ISF. Driven by the influx of CSF, the ISF flushes the brain parenchyma, supplying nutrients [7] while taking away harmful metabolites [8], and finally returns to the subarachnoid space through perivenous routes [9]. Dysfunction of ISF bulk flow would, thus, lead to inefficient cerebral waste clearance, which is linked with many critical brain disorders, such as Alzheimer’s disease, Parkinson’s disease, diabetes, cerebral oedema, and stroke [3, 10]. Dysfunctions of brain metabolites clearance increase with age [3, 11–13], and the associated diseases have been top concerns in brain-disease research and treatments. Some researchers argue that molecules’ thermal motion (diffusion) instead of ISF bulk flow (convection) dominates the waste clearance [14–17], while opponents contended that tracer movement in the brain tissue is much faster than driven by diffusion alone [18–21], which means that that interstitial velocity is fast enough to transport large, slow-to-diffuse molecules [22]. This has been a debate lasting for nearly a decade [23, 24].

CSF and ISF also serve as the drug delivery pathway in the treatments of neurological diseases [25, 26]. Since the blood-brain barrier (BBB) severely restricts the passage of most conventional drugs because of its impermeability [27], the Convection-Enhanced Delivery (CED) method was therefore proposed to bypass the BBB by infusing drugs directly into brain tissues through catheters [28]. Despite being regarded as a promising treatment in the past 20 years as it is broadly applicable to deliver various therapeutic compounds for a diversity of diseases [29, 30], achieving optimal drug distribution in clinical trials remains a challenge [29, 31–33].

## Biomechanics associated with brain interstitial transport and current challenges

Morphologically, the brain resembles porous media; this similarity has inspired engineers and mathematicians to create computational models to deepen our understanding of fluid transport in the brain [34, 35]. Compared to experimental methods, computational modelling can surpass resolution limitations and decouple the complex multiphysics process.

Consequently, models have been established across the scales to predict solute distribution at the brain scale [36–38], analyse fluid transport at the cell scale [15, 16, 39], and study molecule diffusion in the brain interstitium [40]. Furthermore, the introduction of poroelastic theory makes it possible, although at the tissue scale, to consider neuron deformation during the fluid transport process [41–43].

Despite extensive research demonstrating the potential of biomechanical modelling to capture interstitial transport in the brain across various scales, a comprehensive and fully validated framework for accurately visualising *in vivo* interstitial transport remains lacking. Biomechanical studies reveal that neurons are remarkably soft and flexible [44, 45], suggesting that the fluid transport pathways, which is defined by the gaps between neurons, may undergo dynamic deformation during the movement of interstitial fluids [46]. These findings challenge the adequacy of the current convection-diffusion paradigm to fully describe brain transport mechanisms, which may underlie the persistent debate and the inability to reliably predict drug distribution in the CED trials.

## Application of experimentally informed bottom-up modelling

This research aims to develop a new multiscale and multiphysics computational framework underpinned by experiments carried out at different scales to provide accurate predictions of interstitial transport in the ovine brain, which shares significant anatomical and functional similarities with the human brain. By using this framework, one can: (i) understand the fluid-brain interactions from the neuron scale to the organ scale; (ii) precisely visualise fluid transport in the whole brain based on medical images, thus being able to predict drug delivery and disease spread; (iii) optimise brain drug delivery procedures to achieve desired drug distribution in the brain.

To this end, we looked into the fluid dynamics in the ovine brain across the scales. Given that white matter (WM) is generally regarded as the primary site for the bulk flow of ISF [47–50], we focus on WM in this study. Accordingly, unless otherwise specified, “whole brain” refers to the WM of the entire brain in this paper. As shown in Fig 1, we first collected the cytoarchitecture information of ovine brains’ WM using Focused Ion Beam Scanning Electron Microscopy (FIB-SEM) technique and reconstructed representative 3D microstructures of the Corpus Callosum (CC), Corona Radiata (CR) and Fornix (FO), which represent the three major WM regions. Second, based on the 3D microstructures, we developed a finite element model to capture the fluid flow between axons and the corresponding axonal deformation. The simulation results perfectly unravelled the complex interaction between fluid flow and its pathway. We further developed deformation and region-dependent permeability tensors of the ovine brain based on the simulation data. A set of *ex vivo* infusion experiments with fresh ovine WM samples successfully verified the proposed permeability tensor. Finally, we integrated the deformation and region-dependent permeability tensors into the anisotropic and heterogeneous brain models and successfully established the most comprehensive computational framework ever reported for modelling interstitial transport in the brain. To validate its accuracy, we conducted 6 groups of *in vivo* infusion experiments with sheep. We tracked the time-varying distribution of Magnetic Resonance Imaging (MRI) tracer in the sheep brains. We reconstructed the sheep brains’ realistic geometries using MRI images. We registered the localised anisotropy and heterogeneity of the brains based on Diffusion Tensor Imaging (DTI). Results showed that the predicted results agree very well with the *in vivo* experiments. Using this computational model, we further investigated the effects of infusion parameters on the MRI tracer transport in WM, providing valuable insights for improving the efficacy of CED.

**Fig. 1:**
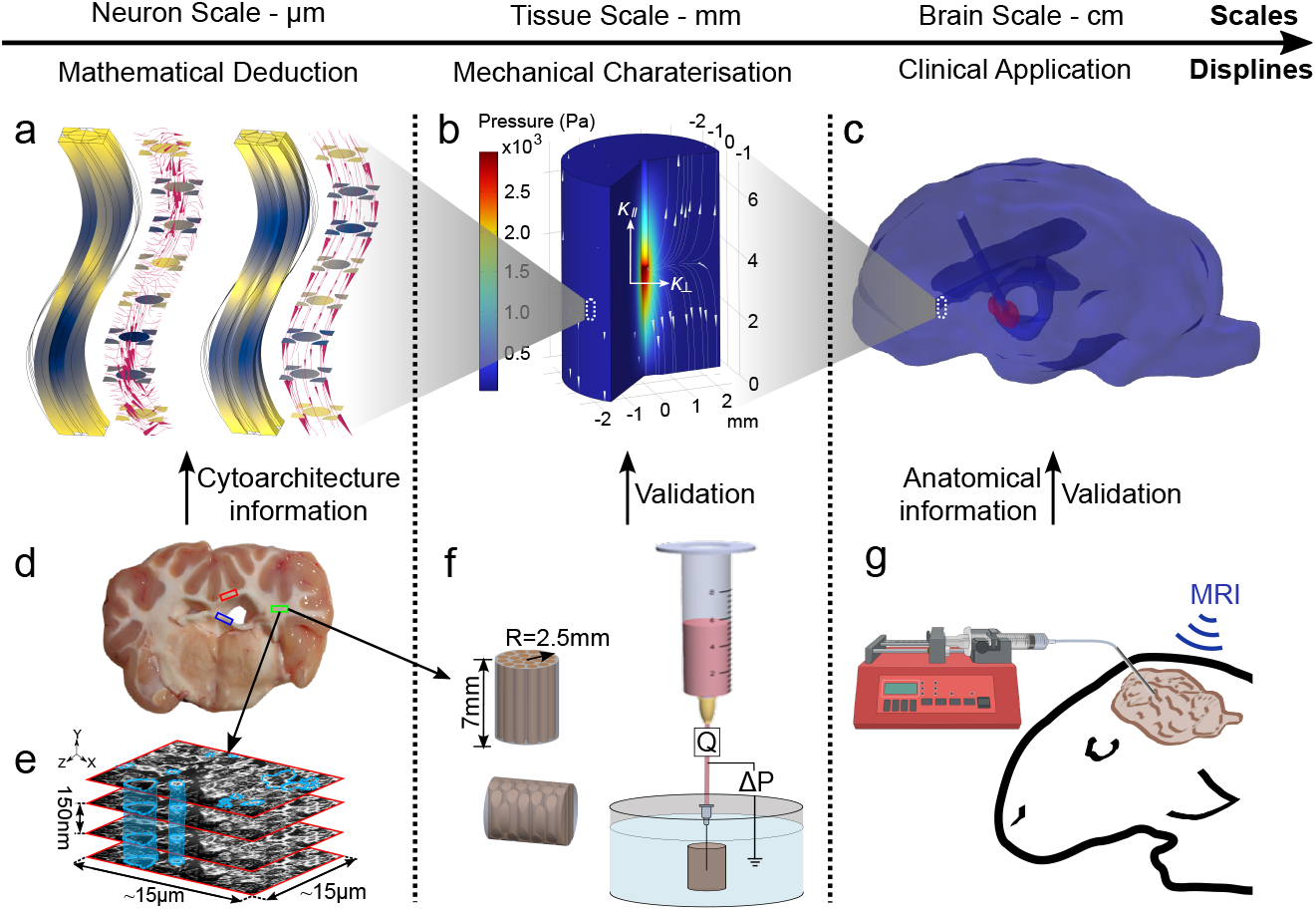
Bottom-up workflow from understanding neuron-ISF interaction to predicting fluid flow in the whole brain. **a**, Fluid-solid interaction modelling demonstrates explicitly how fluid flows through the extracellular space and how axons respond in return. A mathematical relationship between interstitial pressure and the permeability tensor of WM was deduced from the simulation results. The simulations were conducted in representative volume elements (RVEs) that were reconstructed from the cytoarchitecture of fresh ovine brain WM (**d**) obtained by FIB-SEM scanning (**e**). **b**, Up-scaling the neuron scale pressure-dependent permeability tensor to tissue scale. This enabled the validation of the permeability tensor by tissue scale infusion experiments as shown in **f**. By assigning the permeability tensor to fresh brain WM samples, simulations can accurately reproduce the relationship between the infusion pressure and the equivalent permeability tensor of the brain samples, which validated the permeability tensor of ovine brain tissues. **c**, The whole ovine brain model reconstructed from T1-weighted MRI images. The tissue scale permeability tensors were assigned to the computational domain based on the DTI registration. **g**, *In vivo* ovine brain infusion experiments enabled the establishment and validation of the modelling platform for whole-brain intestinal transport. Details of each figure will be given in the main text.

Through this work, we have (i) uncovered the significant effects of axonal mechanical behaviours on fluid transport in WM, challenging and enhancing the existing convection-diffusion paradigm and providing new insights into mechanisms governing interstitial transport in the brain; (ii) developed the first fully-validated multiscale and multiphysics modelling framework in the open literature that can precisely predict *in vivo* solute distribution after brain infusion. We anticipate that our modelling framework will not only improve the efficacy of localized drug delivery strategies, such as CED, for a variety of diseases, but also provide critical tools for the development of novel therapeutic approaches for numerous brain disorders. Furthermore, this framework has the potential to advance our understanding of brain mechanisms, paving the way for new avenues of exploration in neuroscience. This wide-reaching impact highlights the transformative potential of our framework in advancing both clinical treatments and fundamental brain research. A more detailed discussion of the broad impacts of this research is provided in the *Discussion* section.

## Cell scale model: fluid-axon interactions

### The microstructure model

Fig. 2a is the magnetic resonance (MR) diffusion tractography of ovine brain WM. By tracking the diffusion of water molecules around nerve fibres, diffusion tractography can visualise the orientation of axon tracts. It shows that axons appear “wavy” under in situ length conditions [52], as shown in Fig. 2b, which is the real structure of axons obtained by FIB-SEM [51]. However, the resolution of the state-of-the-art MR technology is insufficient to support reconstructing the tissue microstructure. We, therefore, adopted the FIB-SEM technique to scan the tissue and obtained the cytostructure of ovine brain WM in the CR, FO, and CC regions, which are the three major parts of the brain WM [53]. The probability densities of their diameters were measured and presented in Fig. S1a. Based on mean values of the axonal diameter and the typical shape of axons, we built 3D representative volume elements (RVEs) to represent the microstructures of CR, FO, and CC, respectively, as shown in Fig. S1b. Except for the diameter and shape of the axons, other geometric properties of the axons, including tortuosity (*τ*, the ratio of the axon’s arch length and the distance to its two ends; the value of CC, CR, and FO are 1.145 ± 0.141, 0.107, 0.107 ± 0.085, and 1.161 ± 0.117, respectively [53]), distances between axons (0.02 *µ*m to 0.1 *µ*m [54–56]), and porosity (*ϕ*, the volume fraction of the extracellular space, 0.18 ~ 0.3 [57]), were also considered to reconstruct the RVEs. See *Methods* for more details of the microstructural reconstruction of the ovine brain WM.

**Fig. 2:**
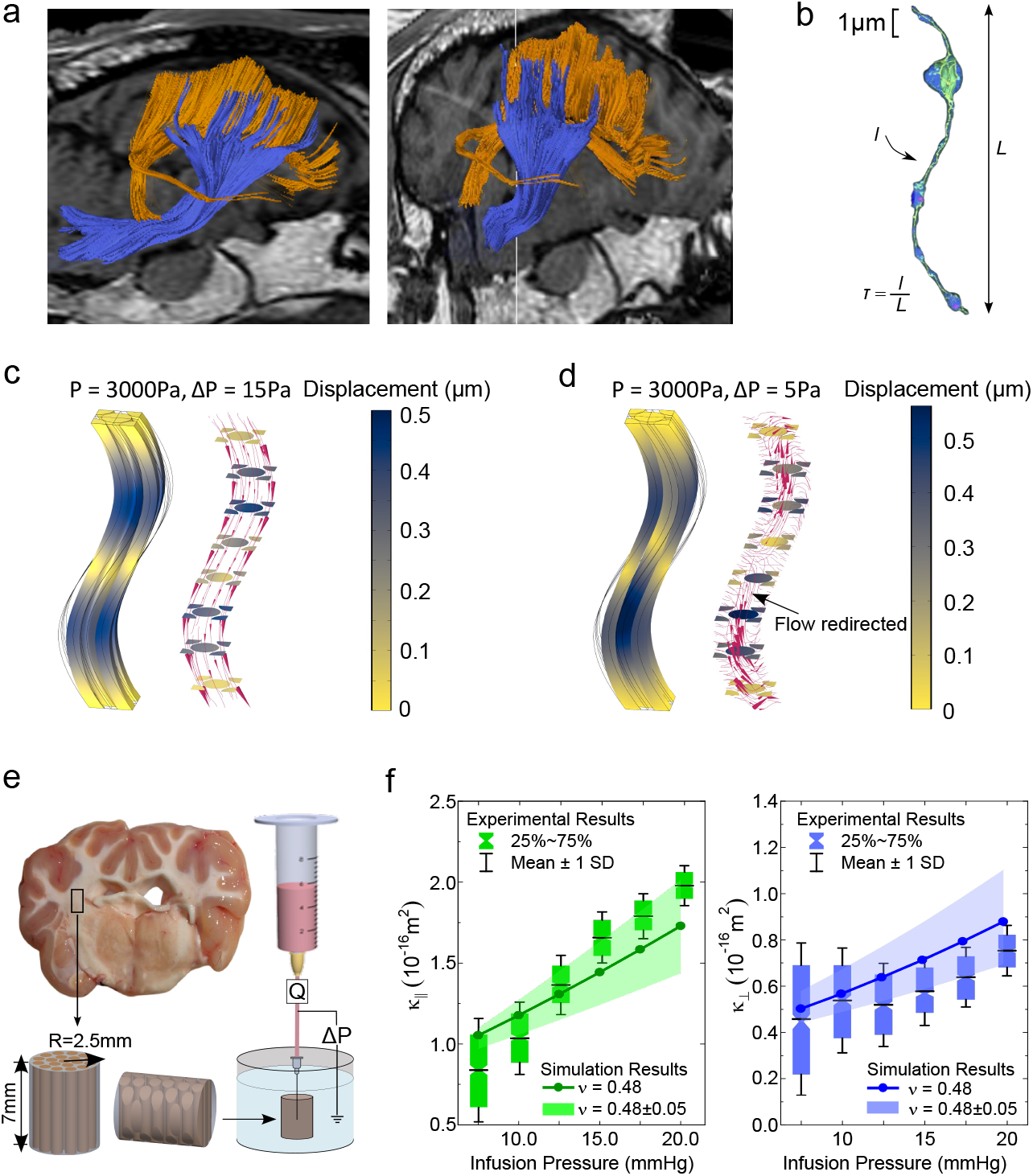
Cell scale and tissue scale models. **a**, The magnetic resonance diffusion tractography of ovine brain white matter. It shows the orientation of axon tracts in CC (orange) and CR (blue) regions. **b**, Real structure of axons obtained by FIB-SEM; reproduced from [51] under open-access terms. **c**, ISF-axons interactions when fluid flows parallel to the axons. The left-hand side figure shows the axons’ deformation (the wireframe shows the original/undeformed shape of the axons) while the right-hand size figure aims to demonstrate the flow status in the deformed flow pathway (the arrows indicate the flow direction and their size is proportional to the flow velocity). **d**, ISF-axons interactions when fluid flows perpendicular to the axons. The interpretation methods are the same as that in **c. e**, Schematic of the *ex vivo* infusion experiment. Tissue samples with nerve fibres in both directions were obtained from the CR region. Infusion pressure ranged from 7.5 to 20 mmHg. The flow rate (Q) and pressure drop along the tissue (P) were monitored to calculate the equivalent permeability of the tissue samples. **f**, Assign the permeability tensor developed above to the tissue sample geometry to model the infusion experiment and then compare the simulation results against the tested results in both directions. The box chart shows the testing results while the lines show the simulation results. Different Possion’s ratios of the axons were tested in the simulations to find the reasonable Possion’s ratio of the axons.

### ISF flow can easily deform axons

Then, we used the reconstructed microstructural RVEs to model the fluid-solid interaction (FSI) between the ISF and axons. As WM is widely treated as transversely isotropic, i.e. cross-sectionally isotropic and longitudinally anisotropic [58–60], two directions of ISF flow, namely parallel to the axons and perpendicular to the axons, were studied. The results are shown in Fig. 2c (parallel flow) and 2d (perpendicular flow).

The normal ICP of adults is 1000 ~ 2000 Pa in a horizontal position and about 252.7 Pa in a vertical position [61]. It can increase to higher than 25 mmHg (3332.5 Pa) under the condition of brain oedema [62]. In CED treatments, the ICP at the infusion site can reach 30 mmHg (3999 Pa) [28]. Therefore, to ensure wide representativity, we chose 0 ~ 3000 Pa as the range of hydraulic pressure in this study, and the results of 3000 Pa were presented in Fig. 2c and 2d. Fig. S1b and S1c also show the results of 0.1 Pa and 1600 Pa in different flow directions. It is worth mentioning that, in real situations, when a different pressure is applied, the pressure gradient will change accordingly depending on the transport properties of the domain. For simplification, we directly adopted the properties of the brain tissue sample used in our *ex vivo* experiments (will be introduced later) to determine the pressure-pressure drop pairs. Under each pair of pressure and pressure drop, we measured the deformation of axons (contours in the left-hand side panels) and the flow status of the ISF (flow stream in the right-hand panels). Results show that, in general, the axons deform under normal ICP. For example, under 1600 Pa pressure, the deformation reaches 0.3 *µ*m (Fig. S1b and S1c), which is approximately 30% of the axons’ average diameter. Under 3000 Pa pressure, their deformations can even reach half of their average diameter.

### Axonal deformation significantly changes the ISF flow

When the ISF flow deforms the axons, it also changes its own pathway. The streamlines in the right-hand side panels of Fig. 2c, 2d, S1c and S1d show how ISF flow status changes with the infusion pressure. The arrows point to the flow directions and their sizes are proportional to the magnitudes of the flow velocity. Under parallel infusion, higher inlet pressure leads to higher flow velocity and more uniformly distributed streamlines. The main reasons are: (1) axonal shrinkage releases more spaces for the ISF flow, as shown in Fig. S2a; (2) flatter pathways due to smaller axonal tortuosity reduce the longitudinal resistance of the ISF flow. Under perpendicular infusion, the difference made by higher pressure is more drastic. Under a higher pressure, the increased pressure gradient partially closes the outlet of the fluid flow [63], as shown in Fig. S2b, causing the majority of streamlines to turn to the parallel direction, as shown in Fig. 2d. This explains the counterintuitive result observed in the *in vivo* brain infusion experiment, where the gadolinium-based bolus, despite being infused perpendicular to the nerve fibres, aligns parallel to them (Figs. 3d, 3g), contrary to the expected perpendicular orientation.

**Fig. 3:**
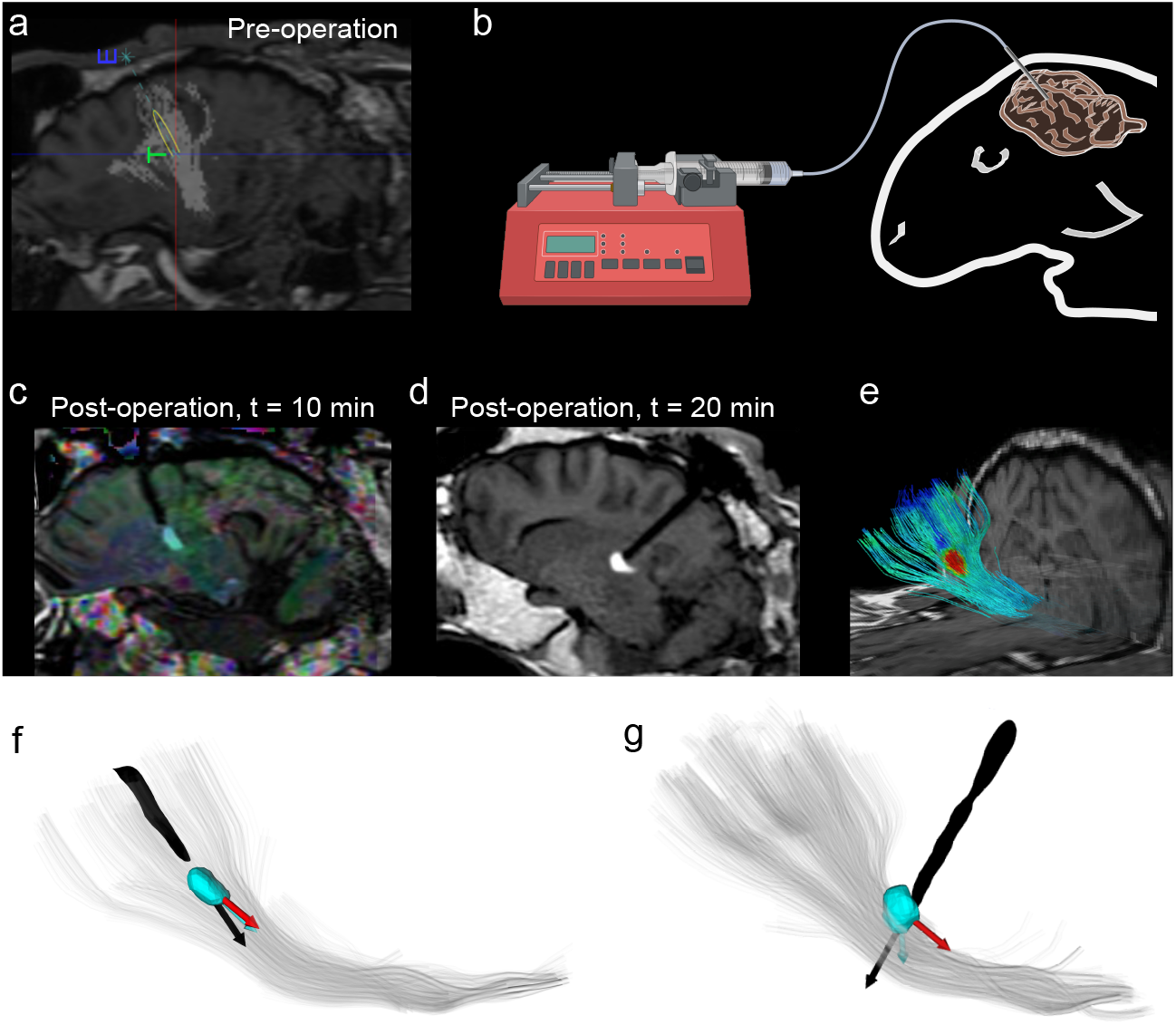
The *in vivo* infusion experiment. **a**, Catheter locating based on diffusion tensor tractography. “E” represents the catheter entering point, and “T” represents the catheter targeting point. The white shadow indicates the orientation of the local nerve fibres. **b**, Infuse the Gadoteridol (Gd) into the sheep’s brain. **c**, Gd infusion parallel to the local nerve fibres. **d**, Gd infusion perpendicular to the local nerve fibres. **e**, Visualisation of the infused Gd and the local nerve fibres. **f**, The relative position of the catheter, the infused Gd, and nerve fibres in the parallel infusion scenario. **g**, The relative position of the catheter, the infused Gd, and nerve fibres in the perpendicular infusion scenario.

## Tissue scale model: pressure-dependent permeability tensor of brain WM

### Theoretical deviation of the permeability tensor

From the macroscopic point of view, the deformation of the ISF flow pathway changes the transport property (which can be characterised by hydraulic permeability ***κ***) of the WM, as our recent work has demonstrated that WM’s hydraulic permeability is highly dependent on its microstructure [35]. Fluids can more easily pass through a porous medium with a higher hydraulic permeability. Note that the hydraulic permeability of WM is transversely anisotropic, so we treated ***κ*** here as a tensor that contains the parallel permeability component and perpendicular permeability component.

According to Darcy’s Law (*κ* = *QµL/A*Δ*P*), where *Q* is the flow rate through the sample, *µ* is the dynamic viscosity of fluid, *L* is the length of the sample, *A* is the cross-sectional area of the domain, Δ*P* is the pressure drop over the length of the domain, the FSI modelling results can provide sufficient information to calculate the localised tissue ***κ*** and relate it to the local hydraulic pressure and pressure drop. Therefore, we set pressure (*P*) and pressure drop (Δ*P*) as parameters to conduct parametric studies and obtained *p* − Δ*p* − ***κ*** relationships in both directions. By fitting curves and surfaces, as shown in Fig. S3, we formulated the relationships as [63]:

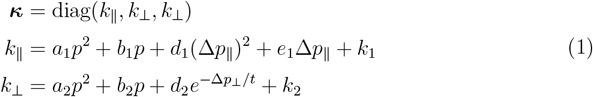

where ***κ*** is the permeability tensor of the brain WM, and the components are *κ*_⊥_ = *κ*_*xx*_ = *κ*_*yy*_ (permeability perpendicular to the nerve fibres) and *κ*_∥_ = *κ*_*zz*_ (permeability parallel to the nerve fibres) according to the transversely isotropic treatment. *a*_*i*_ *b*_*i*_ *d*_*i*_(*i* = 1, 2), *e*_1_, *t* are the coefficients and [*k*_1_, *d*_2_ +*k*_2_] is the permeability tensor without axonal deformation.

More details about the derivation are given in *Methods* and Fig. S3. This newly developed permeability tensor formulation offers more accurate predictions of brain interstitial transport than existing methods, as it quantifies varying transport properties and links them to the local hydraulic status of the tissue.

### Tissue experiments verify the permeability tensor

We conducted a set of *ex vivo* experiments with ovine brain WM to verify the relationship between hydraulic status and tissue permeability tensor, as schematically shown in Fig. 2e [64]. The samples were cylindrical with a diameter of 5 mm and a length of 7mm from the CR zone of WM. Approximately half of the samples have their axes along the nerve fibres while the rest have their axes perpendicular to the axon fibres; the aim was to obtain the two components of the permeability tensor. We inserted a syringe needle (BD MicrolanceTM; stainless steel; 30G × 1/2”; 0.3 ×13 mm) till the middle of the samples and infused PBS into the tissue. According to Darcy’s Law, we measured the pressure drop (P) and flow rate (Q) during the infusion process to calculate the tissue permeability. We applied a range of infusion pressure and drew the permeability tensor against the infusion pressure, as shown by the box charts in Fig. 2f.

We then reconstructed the sample geometry and assigned the newly developed permeability tensor to it to simulate the experiments (see *Methods* for details). The line charts in Fig. 2f present the simulation results. While axons are widely treated as a nearly incompressible material (Poisson’s ratio is between 0.475 and 0.5), the exact value of their Poisson’s ratio is still unknown. As axonal shrinkage is one of the major deformation modes that change the ISF flow status under different hydraulic pressures (analysed above), and Poisson’s ratio determines the compression behaviour, it requires careful attention when defining Poisson’s ratio of axons in this study. Here, we used Poisson’s ratio *ν* = 0.475, 0.480, and 0.485 to define the axons. The results show that the axon’s Poisson’s ratio indeed has a significant effect on the tissue permeability and its exact value should be around 0.480.

Both experimental evidence and numerical results show that tissue permeability increases with hydraulic pressure. The simulated relationship of tissue permeability tensor against infusion pressure also agrees well with that from experimental tests. Therefore, we have validated the FSI modelling method and the derived permeability tensor.

### Permeability tensor of WM in different regions

Despite having unveiled the pressure dependency of WM’s permeability tensor and the mechanism behind the phenomenon, whether different regions and physiological activities may lead to different pressure-permeability relationships is still unknown. We then used the above-validated FSI simulation method to characterise the pressure-permeability relationships of FO and CC. Fig. S4a shows that this relationship in CC and CR is almost the same, while the permeability tensor is more sensitive to infusion pressure in FO. These phenomena may have originated from the microstructure difference [35], as shown in Fig. S1a. Clearly, the axonal diameters in CC and CR are similar but larger in FO. When closely packed, larger axons would lead to larger gaps between axons, thus leaving more spaces for the flow to develop. This explains why the magnitude of permeability is higher in FO. The same goes for the pressure sensitivity of the permeability tensor, i.e., with the same bulk modulus and hydraulic pressure, larger axons also lead to a larger volume shrinkage of axons and more spaces for the flow. This may provide clues to some specific functional differences between FO and CC/CR. For example, FO is a significant bilateral association pathway within the limbic system and also acts as a commissural pathway, connecting the twin parts of the hippocampus in the two hemispheres [65]. This dual role makes FO a crucial component in the brain’s connectivity and function [66, 67]. The high requirement of interconnection should require FO to have a higher capability of transferring matters.

## Organ scale model: a whole brain transport modelling framework

### *In vivo* brain infusion experiments

With the permeability tensors of the brain WM in different regions, we could obtain the distribution of fluid transport property in the whole brain WM. This enabled the establishment of the organ scale model. To validate the accuracy of the organ scale model in predicting the whole brain fluid transport process, we have conducted 6 groups of *in vivo* infusion experiments with sheep.

Guided by MR imaging, we inserted a hollow guide catheter into the internal capsule (associated with the CR region) of the ovine brains while making the guide catheter parallel (groups 1 ~ 3) and perpendicular (groups 4 ~ 6) to the axon tracts, as shown in Fig. 3a. We then infused gadolinium-based solution (Prohance^®^, 1:80 in saline, MRI tracer) via a specialised silica catheter that was inserted directly into the target through the hollow guide-catheter (Fig. 3b) and tracked the time-varying gadolinium distribution by 3D T1-weighted Fast Field Echo (FFE) (Fig. 3c and 3d), at multiple time points. Binary masks of the infused boluses for all time points were extracted by segmentation of the Gadolinium distribution on post-injection 3DT1 images. We also extracted the corticospinal tracts from the whole brain tractography (Fig. 3e) and calculated the principal directions (eigenvectors, *ε*_1_, *ε*_2_, *ε*_3_), principal length (*λ*_1_, eigenvalue of *ε*_1_), and volume of each bolus at different time points. After spatial alignment, we finally registered the bolus into the fibre tracts (Fig. 3f and 3g) to understand how fibre direction, or flow direction, affects the interstitial fluid flow and substances transport. See *Methods* for more details.

It was observed that, while the principal direction of the Gd bolus infused parallel to the fibre tracts remains aligned with the fibres (Figs. 3c, 3f), the principal direction of the drug bolus infused perpendicular to the fibre tracts does not align as expected (Figs. 3d, 3g). This discrepancy is fully explained by the fluid-axons interaction simulations, which reveal that the perpendicular infusion partially obstructs the outlet, leading to a redirection of the flow in the parallel direction (Fig. 2d).

### The computational model

Before infusion experiments, we scanned each sheep’s brain using MRI. The T1 weighted MR images were then used to reconstruct the whole brain geometry, as shown in Fig. 4a. This provides the geometry needed to develop the modelling framework for the whole brain transport. Considering the anisotropy of WM, we also extracted the diffusion tensor of the water molecules with the voxel resolution of 0.2 × 0.2 × 0.2 *µm*^3^, as shown in Fig. 4b, based on which the direction of nerve fibres can be obtained. The zoomed-in part illustrates the principal directions of the local water molecule diffusion tensor; the component with the largest magnitude in each voxel points to the local direction of axon fibres. We then assigned the permeability tensor developed above to each voxel, with *k*_∥_ aligned to the axonal direction. The diffusion coefficient of Prohance was adopted after orthogonalisation based on the diffusion tensor in each voxel [68]. Fig. 4c shows the geometry of a whole ovine brain with permeability tensor and diffusion tensor information registered. Mathematically, we employed the convection-diffusion-reaction equation to model fluid flow and substance transport in the brain; see *Methods* for the details. Fig. 5a and 5b show the results of a parallel infusion case and a perpendicular infusion case, respectively.

**Fig. 4:**
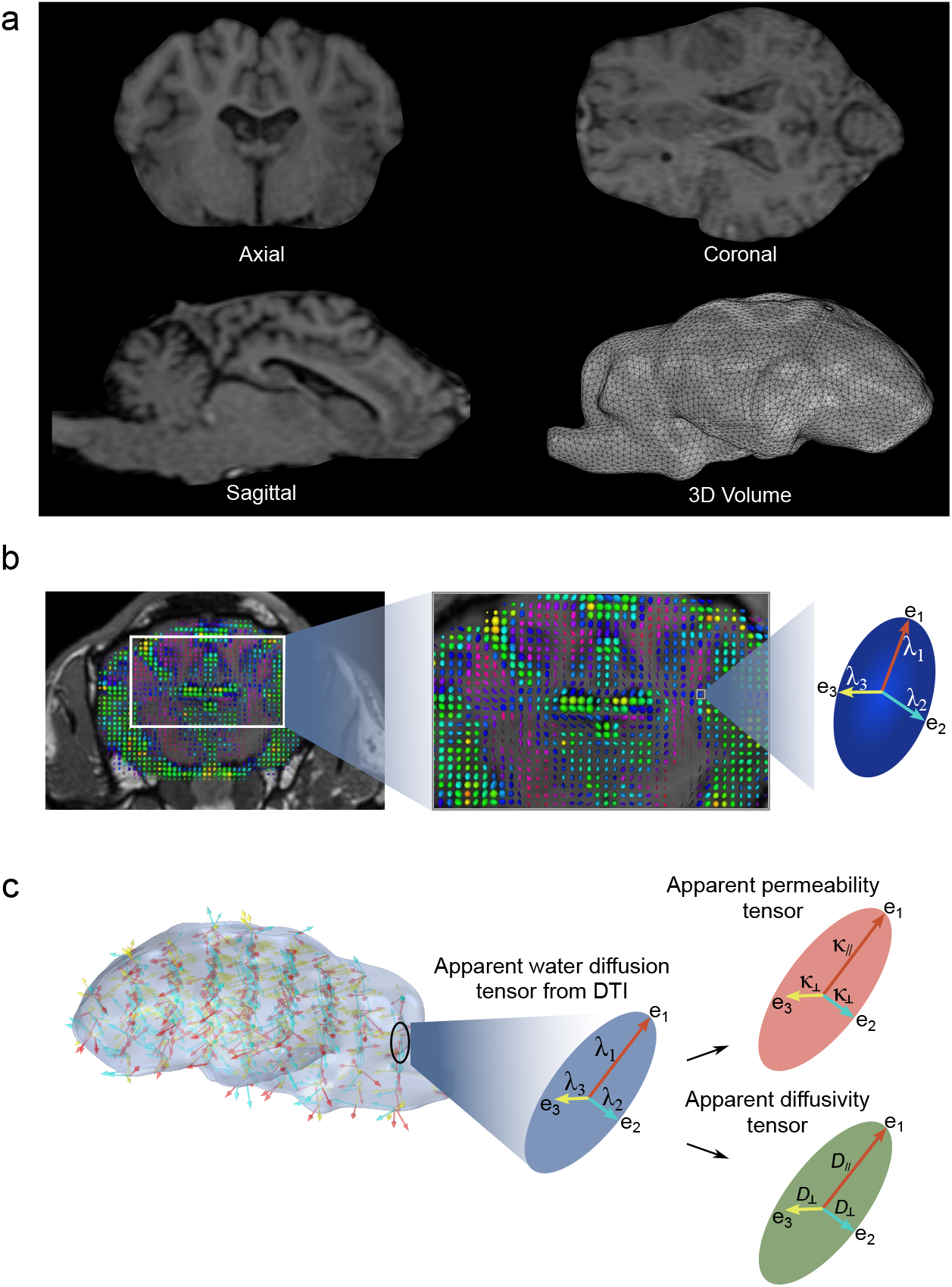
Brain transport modelling framework. **a**, T1 weighted MR image of each sheep for brain geometry reconstruction. The 3D volume shows the finite element meshed brain geometry. The mesh around the infusion site was refined to balance high accuracy and affordable computational burden. **b**, Visualisation of the local diffusion anisotropy of water molecules. The long axis of the ellipsoid aligns with the major diffusion direction of water molecules, indicating the local orientation of nerve fibres. This information can be used to register the local anisotropy in the brain geometry. **c**, Diffusion Tensor Imaging (DTI) data was used to register the local anisotropy to the brain geometry. After the registration of DTI information, the direction of transport properties of each element is aligned to the axon fibres.

**Fig. 5:**
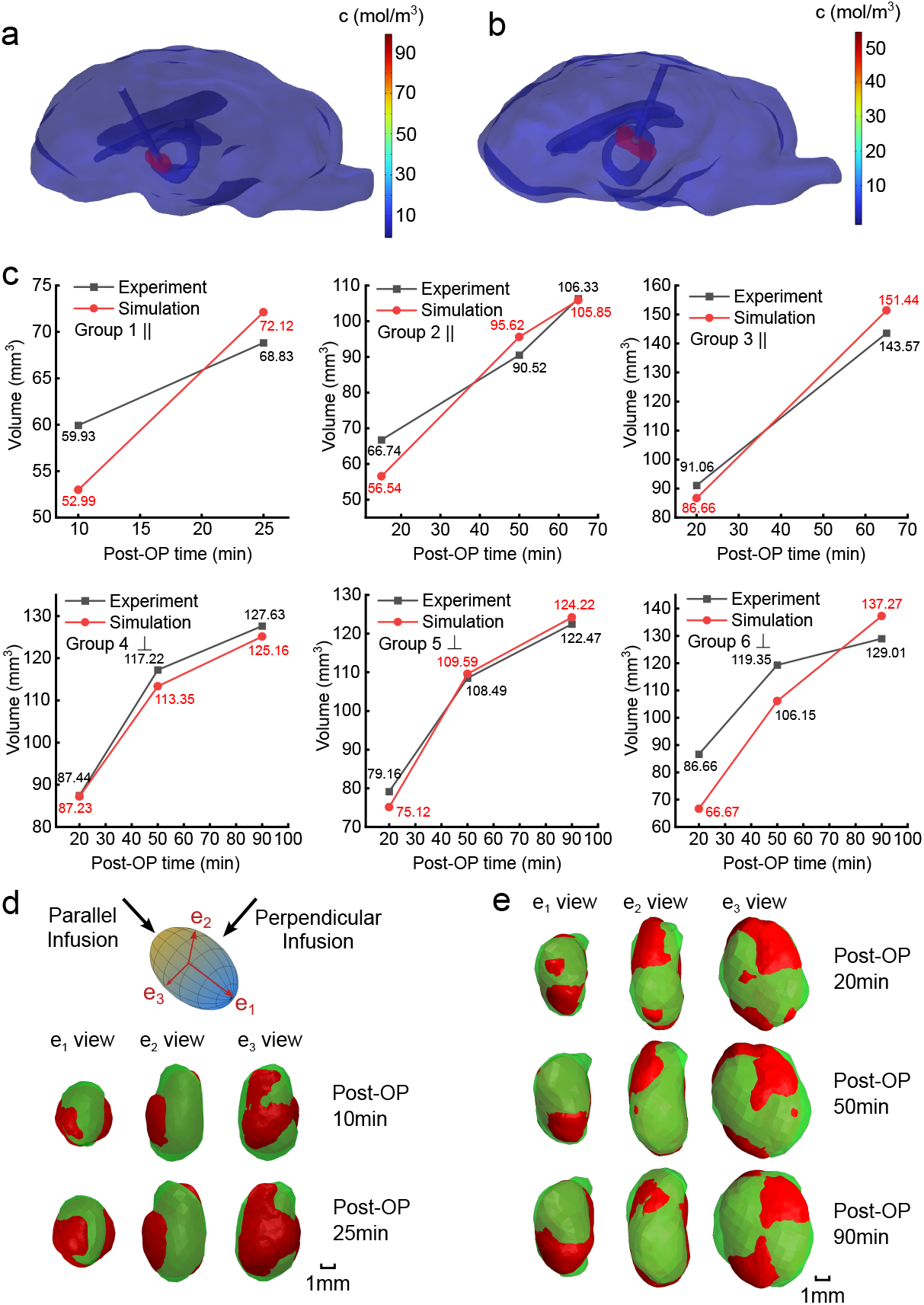
Simulation results and model validation against *in vivo* experiments. **a**, The Gd concentration distribution in the parallel infusion case. The red bolus is the area where the Gd concentration is higher than 0.004mol/L, corresponding to the Gd detection boundary under MRI. **b**, The Gd concentration distribution in the perpendicular infusion case. **c**, Comparison of the time-varying size of Gd bolus after infusion in the 6 groups of sheep brains. **d**, Comparison of the Gd bolus shape obtained by experiment (green) and simulation (red) at different time points after parallel infusion. The results are from group 1. **e**, Comparison of the Gd bolus shape obtained by experiment (green) and simulation (red) at different time points after perpendicular infusion. The results are from group 4.

### Model validation

To validate the computational model, we compared the shape and size of the Gd boluses obtained from simulations (red) and experiments (green) at different infusion time points. The presented bolus volumes represent regions where the Gd concentration is higher than 4 mmol/L. While Fig. 5c compares the sizes of the Gd boluses in all six groups at different time points, Fig. 5d and 5e future compare the shapes of the boluses infused parallel to the nerve fibres (group 1) and perpendicular to the nerve fibres (group 4), respectively, from different perspectives. The shape comparisons of other groups are shown in Fig. S5. The results show that the simulation results agree well with the experimental results, demonstrating the capability of the newly proposed bottom-up modelling framework to predict fluid flow and substance transport in living brains.

## Model application: optimising CED efficacy in the brain

Using the newly developed model, we analysed the impact of infusion parameters on the delivery outcomes of CED, as illustrated in Fig. 6. These findings provide valuable insights for optimising CED efficacy in the brain.

**Fig. 6:**
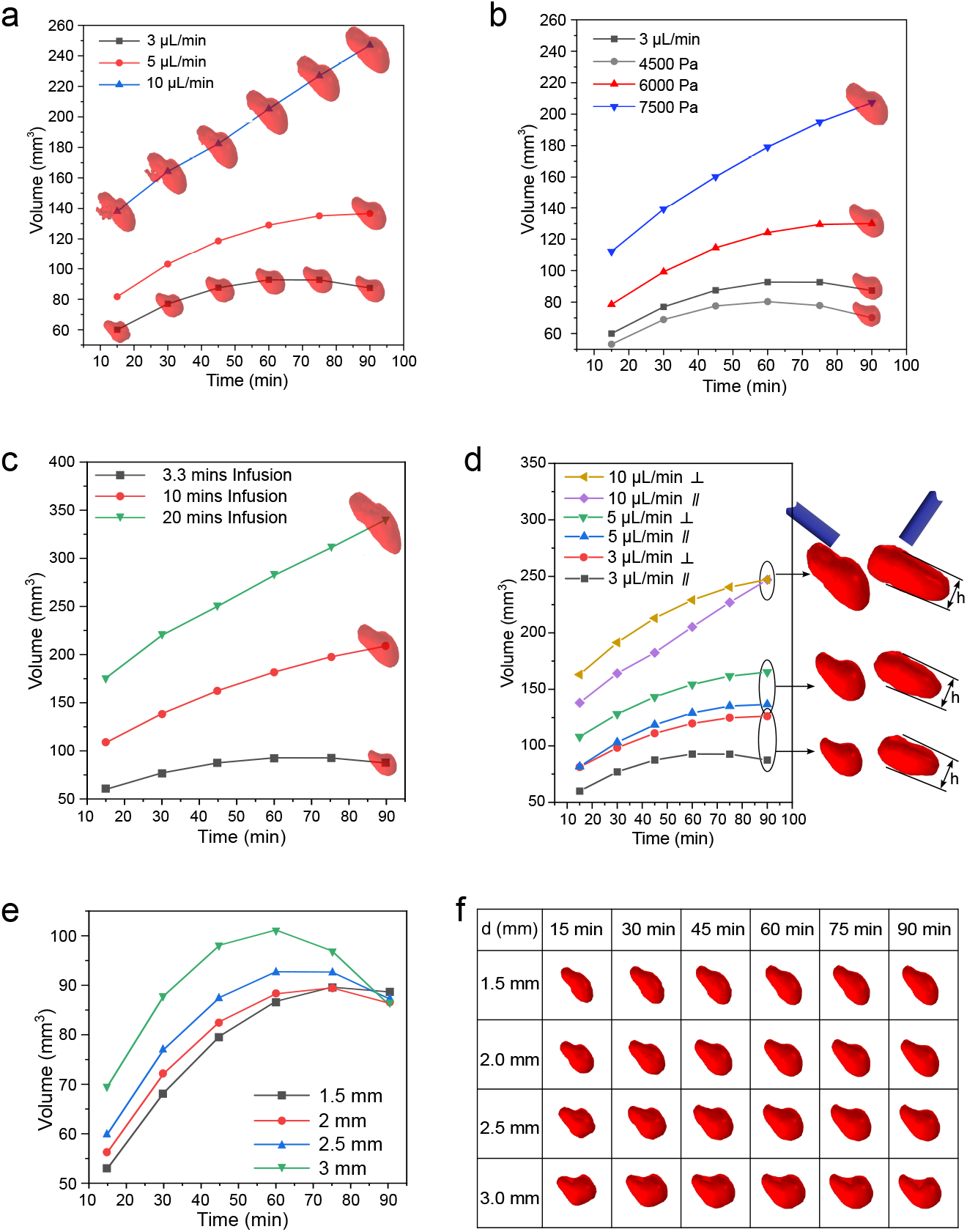
Effects of infusion parameters on the distribution of the Gd bolus. Unless otherwise specified, the infusion duration was 3.3 minutes, the infusion rate was 3 *µ*L/min, and the red boluses represent the regions where the Gd concentration was no lower than 4 mmol/L in all panels. **a**, Effects of infusion rate on the Gd volume in the brain after the end of infusion. **b**, Effects of infusion pressure on the Gd volume after the start of infusion. **c**, Effects of infusion duration on the Gd volume in the brain after the start of infusion. **d**, Effects of infusion direction on the Gd volume in the brain after the start of infusion. **e, f**, Effects of catheter size on the Gd volume and shape, respectively, in the brain after the start of infusion.

### Effects of the infusion rate and pressure

Three infusion rates (3 *µ*L/min, 5 *µ*L/min, and 10 *µ*L/min) were tested to evaluate their effects. At a flow rate of 3 *µ*L/min, the infusion pressure reached 4500 Pa. To further investigate the effects of pressure-controlled infusion, we examined delivery outcomes at pressures of 4500 Pa, 6000 Pa, and 7500 Pa, comparing these results with those of flow rate-controlled infusion. For simplicity, only the parallel infusion direction was considered.

As shown in Fig. 6a and 6b, the bolus volume increases with both infusion rate and pressure. This occurs because a higher infusion rate or pressure delivers more Gd into the brain within the same time. At 3 *µ*L/min (or 4500 Pa), the bolus volume starts decreasing approximately 1 hour after infusion. However, this trend gradually reverses as the infusion rate or pressure increases. At 10 *µ*L/min (or 7500 Pa), no decrease in bolus volume is observed within 1.5 hours post-infusion. These results suggest that a low infusion rate may fail to maintain effective drug concentrations, as it delivers an insufficient dose, further exacerbated by rapid dispersion due to convection-driven transport. The short infusion duration (3.3 minutes) also contributes to this issue. To address this, longer infusion durations were investigated in the next Section.

Results also indicate that higher infusion rates or pressures produce a more anisotropic Gd distribution. For instance, at 90 minutes, the bolus from a 10 *µ*L/min infusion exhibits a longer major axis but a similar minor axis compared to the bolus at 3 *µ*L/min. This aligns with findings from the abovementioned cell-scale model, where permeability parallel to axons increases more rapidly with pressure than perpendicular permeability.

Notably, although infusion rate and pressure produce similar drug distribution patterns, pressure-controlled infusion (e.g., 4500 Pa, equivalent to 3 *µ*L/min) results in smaller boluses compared to flow rate-controlled infusion (Fig. 6b). Thus, flow rate control is more effective for maintaining efficient drug concentrations. While higher pressures can achieve similar results, they increase the risks of tissue damage and backflow.

### Effects of the infusion duration

To investigate the effects of infusion duration, two additional durations (10 minutes and 20 minutes) were applied, in line with clinically used ranges [69, 70]. Since these durations are comparable to the monitoring period (90 minutes), the Gd boluses were tracked from the start of infusion. As shown in Fig. 6c, longer infusion durations sustain effective drug concentrations for a longer time, as shown by the increased volume of regions where the Gd concentration remains above 4 mmol/L at all time points. This confirms that the decrease in drug concentration after 1.5 hours at a 3 *µ*L/min infusion rate is not solely due to the low infusion rate; the short infusion duration also plays a critical role. An optimised combination of infusion rate and duration is necessary to maintain effective drug concentrations within the target region.

Results also show that longer infusion durations lead to more anisotropic drug distribution. The mechanism aligns with the effect of higher infusion rates on anisotropy, as explained in the previous section.

### Effects of the infusion direction

The influence of infusion angle on drug distribution was also examined. Based on spatial and angular data from the *in vivo* experiment, Gd delivery outcomes were simulated for infusions parallel and perpendicular to local nerve fibres under different flow rates (Fig. 6)d. While both directions exhibit similar growth trends for the Gd bolus, notable differences in volume were observed.

At lower infusion rates, Gd boluses in the perpendicular direction grow faster than those in the parallel direction, indicating that perpendicular infusion can sustain effective drug concentrations for longer. However, this difference diminishes with increasing infusion rates, and at 10 *µ*L/min, the trend reverses after 1.5 hours. In parallel infusion cases, Gd transport is accelerated by convection, leading to a faster decrease in *in situ* concentration and slower bolus growth. Conversely, in perpendicular infusion, convection-driven transport along the catheter direction is mitigated by the resistance of horizontally aligned nerve fibres.

This difference is obvious at low infusion rates, where fluid-axon interaction remains limited. At higher infusion rates, lateral displacement of axons caused by the infused drug becomes more pronounced. In perpendicular infusion cases, the increasing flow rate/pressure gradually closes gaps between nerve fibres. Once these gaps are fully closed, drug transport becomes confined to the direction of the nerve fibres, as reflected in the limited growth of the minor axis of perpendicular boluses with higher infusion rates.

It is worth noting that while perpendicular infusion at low flow rates sustains effective drug concentrations for longer, the penetration depth (major axis length) is shorter compared to parallel infusion.

### Effects of the catheter size

Catheters with diameters of 1.5 mm, 2 mm, 2.5 mm (as used in the experiment), and 3 mm were tested to investigate the effect of catheter size on Gd distribution. In all cases, the infusion rate was 3 *µ*L/min, and the infusion duration was 3.3 minutes. As shown in Fig. 6e, larger-diameter catheters delivered drugs at effective concentrations over a larger region under identical infusion conditions.

An analysis of bolus shapes in Fig. 6f reveals the underlying mechanism: larger catheters increase the tissue area exposed to the drug and reduce the anisotropy of drug distribution. However, the volume of the bolus decreases more rapidly when using larger catheters, resulting in a shorter duration of effective drug concentration. Additionally, larger catheters cause more significant nerve and tissue damage [71].

In summary, while larger catheters achieve more isotropic drug distribution in anisotropic brain tissue and expand the effective dose area, they also shorten the duration of drug efficacy and increase tissue damage. The reduced duration of efficacy could potentially be mitigated by extending the infusion duration.

## Discussion

### Mass transport in the brain: diffusion driven or convection driven?

The question of whether molecular transport in the brain is driven primarily by advection or diffusion remains a subject of ongoing debate. However, little attention has been given to the role of microstructural heterogeneity in influencing these transport mechanisms. Recent advancements in non-invasive imaging techniques have enabled the tracking of molecular transport and fluid flow within the brain interstitium *in vivo* [4, 10, 14, 17, 72], facilitating major breakthroughs, such as the elucidation of the relationship between sleep and metabolite clearance [8]. Nevertheless, the resolution of these techniques does not support tracking the brain tissues’ microstructural change during the mass transport processes [73].

Our study introduces a novel perspective by demonstrating that molecular advection in the brain can vary significantly across different brain regions and under various physiological conditions, largely due to underlying microstructural differences. As illustrated in Fig. S4, the regional variations of brain tissues’ microstructural properties, along with local hydraulic pressure, exert a substantial impact on tissue permeability, which is a key determinant of convective transport in porous media. We find that factors such as regional tissue composition and porosity can dramatically influence convection strength. For example, the tissue permeability in the FO is more than double that in the CC and the CR. Furthermore, increasing tissue porosity from 0.2 to 0.3 results in a fivefold rise in permeability, with both porosity values commonly observed in different brain regions [74]. These findings suggest that the relative importance of advection versus diffusion in brain transport is highly context-dependent, varying with tissue microstructure, regional characteristics, and local conditions. Therefore, the driving force behind molecular transport in the brain is not solely dictated by a singular mechanism but by a complex interplay of microstructural and micro-environmental factors.

### New insights into how sleep enhances brain waste clearance

The porosity of brain tissue is a dynamic variable, significantly influenced by physiological states such as sleep. Research has demonstrated that the volume fraction of the extracellular space (ECS) can increase by up to 50% during sleep [8]. This finding has been central to the argument that sleep enhances waste clearance in the brain [8, 75]. Our simulation results in Fig. S4c provide further quantitative support for this hypothesis. Specifically, they reveal that a 50% increase in porosity (e.g. from 0.2 to 0.3) results in a fivefold increase in permeability, drastically facilitating the convective transport of interstitial fluid.

Such a marked rise in permeability not only accelerates the clearance of metabolic waste but also enhances the transport of essential substances, including nutrients, oxygen, and therapeutic drugs. This relationship underscores the broader physiological significance of porosity modulation in the brain ECS. Changes in ECS volume fraction during sleep or other physiological activities likely play a critical role in maintaining homeostasis and enabling efficient molecular transport.

### Versatility of the newly developed prediction framework

The newly developed modelling framework for predicting fluid and substances transport (interstitial transport) pathways in the brain offers several key applications, particularly in advancing medical and clinical practices. One of its most promising capabilities is the ability to predict interstitial transport in the brain based on medical images. This feature could play a crucial role in identifying the early stages of brain diseases, such as Alzheimer’s, Parkinson’s, and brain tumours [3, 76], by detecting abnormal transport processes before clinical symptoms appear, potentially enabling early diagnosis. By understanding how diseases affect interstitial transport, the framework also provides critical insights into the mechanisms of disease progression, helping to tailor interventions and therapies more effectively.

In addition, this framework can optimise drug delivery by accurately modelling how therapeutic agents are distributed within the brain, leading to more targeted and effective treatments. For example, it can significantly enhance the efficacy of CED by accurately simulating drug distribution within the brain. As mentioned before, CED faces challenges such as uneven drug distribution due to tissue heterogeneity; the framework’s ability to model drug delivery pathways can help optimise these processes. It can predict the most effective injection sites, flow rates, and drug clearance patterns, improving uniformity and reducing toxicity. The framework also accounts for the dynamic nature of brain tissue, including its varying porosity and stiffness, allowing for more precise targeting in both healthy and diseased brain areas. By modelling the effects of pathological conditions, such as tumours, it can adjust delivery strategies to maintain therapeutic efficacy. In essence, the framework can improve CED’s precision and effectiveness, leading to more targeted, personalised, safe, and effective treatments. Additionally, the framework’s flexibility allows it to support tissue engineering and regenerative medicine applications by guiding the design of novel biomaterials and drugs that integrate more effectively with various tissues, improving therapeutic delivery.

Furthermore, the framework can contribute to the development of brain-machine interfaces, such as syringe-injectable neuro electronics [77]. By accurately predicting the transport dynamics during the implantation, it can help refine the delivery and positioning of devices, potentially overcoming the surgical challenges associated with neuroprobe employments, especially tissue damage. This could enhance the precision and safety of neuro electronic implants, improving their integration and functionality within the brain. Through these applications, the framework has the potential to enhance diagnostics, treatment strategies, and surgical planning, with far-reaching implications in medicine.

Its adaptability to other tissues, such as those involved in cancers, greatly increases its application potential across diverse medical fields.

### Impact of fluid-axon interactions on CED

Compared to existing mathematical models for brain drug delivery, a key improvement of the present framework is the incorporation of fluid-axon interactions at the neuronal scale, which are then upscaled to the macroscale through a permeability tensor that varies with brain region, local pressure, and pressure gradient. This enhancement not only underscores the importance of examining how microscale fluid-axon interactions influence macroscale drug delivery outcomes via the CED technique but also provides theories and practical tools to qualify this influence.

Fig. S6 shows the effects of fluid-axon interactions on the Gd distribution under varying flow rates, infusion durations, and infusion directions. In parallel infusion cases, incorporating these interactions results in faster growth of the Gd bolus due to increased parallel permeability. Conversely, in perpendicular infusion cases, fluid-axon interactions reduce the growth rate of the Gd bolus because the perpendicular permeability decreases as gaps between axons close under specific combinations of local pressure and pressure gradients. A comparison across the panels shows that at low infusion rates and short durations, the effect of fluid-axon interactions remains minimal in both infusion directions. However, the impact of these interactions becomes pronounced with increases in either infusion rate or duration. These findings underscore the critical role of fluid-axon interactions in determining drug distribution patterns in the brain.

Additionally, it is important to note that intracranial pressure changes over time. Such changes can significantly influence interstitial transport, potentially contributing to the development of brain diseases. Understanding these pressure-driven effects is essential for improving drug delivery strategies and disease management.

### The translational relevance enhanced by using ovine brain

The ovine animal model has been selected for this study due to its anatomical and functional similarities to the human brain, enhancing the translational relevance of the research [78]. Unlike murine models, which have lissencephalic (smooth) brains devoid of sulci, the ovine brain is large and gyrencephalic, featuring sulci that more closely resemble human brain morphology [79]. Moreover, ovine cerebral vasculature demonstrates notable similarities to the human brain, sharing comparable characteristics observable under radiological imaging [80] and electroencephalogram monitoring [81], as well as functional properties [82]. Therefore, the new findings and newly developed prediction framework in the present study have great potential to be adapted to the human brain.

As part of the Horizon EDEN 2020 project, the co-authors have developed a modular robotic system for precision neurosurgery featuring a bio-inspired, implantable steerable needle designed for a range of diagnostic and therapeutic applications, with an initial focus on localised drug delivery [83]. Integrating this robotic platform with our predictive engine could enable unprecedented accuracy in brain-targeted drug delivery, maximising precision while minimising invasiveness.

### Limitations and perspectives

Despite its robustness and comprehensiveness, the newly developed prediction framework has limitations that warrant further investigation.

#### Microscale Limitations

As a multiscale framework, the overall accuracy strongly depends on the precision of microscale results and the transfer of information across scales. Our previous studies [40, 84] and Fig. S4 highlight the significant influence of tissue microstructure on convection and diffusion in interstitial transport. In this work, we used statistically reconstructed microstructures rather than directly measured inputs. The prediction accuracy could be improved by integrating more realistic geometric models and developing high-throughput methodologies to extract RVEs directly from 3D brain images.

Furthermore, different brain regions exhibit distinct microstructures and transport properties. While this study explored the CC, CR, and FO within the WM, future work should include other regions, including the grey matter. Unlike the WM, the grey matter comprises neuron cell bodies, leading to unique packaging patterns. Nevertheless, the same methodological approach can be applied. Notably, Amunts et al. [85] have developed a 3D probabilistic atlas of the human brain’s cytoarchitecture, which could significantly enhance this framework for guiding drug delivery strategies.

#### Biochemical Interactions

Due to the stability of Gd as the MRI tracer, the effects of biochemical reaction in the present study was limited. For drug delivery and molecular transport modelling, the reaction terms, which covers e.g. molecular binding and endocytosis, could be critical. Lower-scale simulations, such as molecular dynamics (MD) and coarse-grained molecular dynamics (CGMD) [86, 87], are well-suited to model these detailed particle-cell interactions. Integrating MD and/or CGMD simulations with the current framework would provide more precise insights into drug development and pharmaceutical research.

#### Tissue-Scale Limitations

At the tissue scale, the permeability tensor estimation did not explicitly consider contributions from the extracellular matrix (ECM) and other biophysical or biochemical processes due to their complexity and a lack of experimental techniques for validation at such scales. To address this, we adopted a lumped-system approach [88], where a single global parameter, such as viscosity, approximated the overall effects of these processes. Although this approximation yielded satisfactory results, future work could employ first-principles simulations to examine the influence of specific interactions, particularly for understanding the effects of varying drug formulations targeting different cell types.

#### Brain-Scale Assumptions

At the brain scale, we assumed identical tissue microstructure and Gd elimination rates across different sheep brains due to a lack of subject-specific data. This assumption led to variations in accuracy across the six groups. While patient-specific measurements of microstructure and drug elimination rates remain unrealistic, mathematical modelling based on high-resolution medical imaging could help characterise these parameters, potentially improving predictive accuracy.

#### Further Developments for Broader Applications

The developed framework serves as a versatile platform for fluid and substance transport modelling in the brain and beyond. Further developments for broader applications include but are not limited to: (i) developing governing equations for transvascular drug transport to design novel blood-brain barrier (BBB) crossing strategies; (ii) adding the cerebral blood system to estimate brain bleeding risks and guide treatment strategies; (iii) combining with poroelastic theory [74] to model hydrocephalus progression and backflow during drug injections.

In the future, we aim to integrate this mathematical framework with medical imaging techniques to create a rapid, unified system for early diagnosis of brain diseases related to interstitial transport and precision drug delivery treatments. Additionally, we are extending the framework to support drug delivery in other tissues, such as oesophageal tumours, and exploring its application in developing bio-mimetic synthetic materials, such as functionalised and composite hydrogels.

To conclude, we have developed and fully validated a novel, bottom-up mechanistic framework that integrates advanced mathematical modelling, ultra-high-resolution imaging, and biomechanical experiments to achieve precise *in vivo* predictions of the transport of fluids and substances within the brain parenchyma. Demonstrated through accurate modelling of MRI tracer transport in living sheep brains, this platform provides unprecedented predictive capability for fluid and substances transport in the brain and insights into the interplay of fluid dynamics and tissue properties. This transformative approach has the potential to significantly advance drug delivery strategies in the brain, cancer, and other soft biological systems. By overcoming limitations in modelling complex transport in soft tissues, our work establishes a robust platform with profound implications for biomedical engineering and translational medicine, promising to accelerate the development of targeted therapies and advancing neurosciences.

## Methods

### Microstructural reconstruction based on FIB-SEM

To achieve a comprehensive understanding of the microstructure of WM and reconstruct it with high fidelity for the proposed modelling framework, our group has previously employed the FIB-SEM technique to build a detailed cytoarchitecture database of three regions of ovine brain WM: the CC, CR, and FO [53]. Using a FIB, layers of tissue were sequentially milled at 150 nm intervals, with the SEM capturing the top surface after each milling step. The full cytoarchitecture was reconstructed by integrating these layers. Analysis of the cytoarchitecture revealed that the axon diameter distributions closely follow a lognormal distribution, as shown in Fig. S1a for each region. Additionally, the average tortuosity (*τ*), defined as the ratio of an axon’s arc length to the straightline distance between its endpoints, was determined to be 1.127 ± 0.109 for the CC, 1.10 ± 0.07 for the CR, and 1.149 ± 0.088 for the FO. Tortuosity, a key parameter characterising axonal shape, is shown in Fig. 2b. For statistical analysis and without loss of generality, the cross-sections of nerve fibres were approximated as circular. Based on the geometric characteristics of CR, a simplified representation of the WM microstructure was reconstructed, as illustrated in Fig. S1b, to investigate fluid-axon interactions. This model facilitates explicit simulation of fluid flow both parallel and perpendicular to the axons under the configurations depicted in Fig. S1b. While the actual microstructure of WM is more complex, incorporating excessive geometric details would introduce irreparable numerical divergence, particularly due to highly nonlinear factors such as fluid-structure interaction and multi-body contact. Therefore, after carefully balancing geometric fidelity and numerical feasibility, this simplified geometric model was selected.

### Mathematical modelling of the ISF-axons interactions

The fluid was modelled as an isothermal, incompressible Newtonian fluid governed by the Navier–Stokes (NS) equations. Due to the low Reynolds number, inertial terms were neglected. By also neglecting gravity, the ISF flow was described by Eqs. (2) and (3).

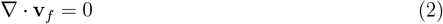

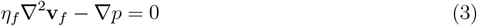

where **v**_*f*_ is the flow velocity, *η*_*f*_ is the dynamic viscosity of ISF (0.8 mPa·s [89]), *p* is the local hydraulic pressure.

The axons were treated as hyper-viscoelastic materials [45, 90]. The total stress in the axons is the sum of the hyperelastic and viscoelastic components [91]:

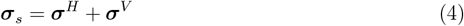

where ***σ***_*s*_ is the total stress of the solid phase (axons), ***σ***^*H*^ is the hyperelastic stress, and ***σ***^*V*^ is the viscoelastic stress.

The hyperelastic response was described using the Ogden model [52], with the strain energy density function defined as:

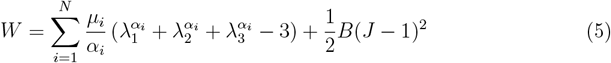

where *W* is the strain-energy function that is expressed in terms of the principal stretches *λ*_*i*_, *i* = 1, 2, 3; *N* is the order of the model, which is 1 for axons; *µ*_*i*_, *α*_*i*_ are material constants with the values of 281.84 Pa, and 6.33, respectively [52]; *B* is the bulk modulus, and 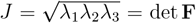 is the volumetric deformation, and **F** = *f* = ∂**x** */*∂**X** is the deformation gradient tensor. The Cauchy stress for the hyperelastic response is:

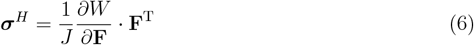

The viscoelastic response, driven by hydraulic pressure and dominated by creep, was modelled using a nonlinear Kelvin–Voigt framework. The viscoelastic stress is given by:

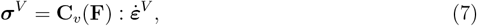

where **C**_*v*_(**F**) is the viscoelastic stiffness tensor, which depends on the deformation gradient **F**, and 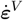 is the rate of the viscoelastic strain tensor. Under sudden stress ***σ***_0_, the nonlinear creep behaviour is:

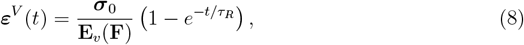

where *t* is time, **E**_*v*_(**F**) is the nonlinear elastic modulus, and *τ*_*R*_ is the retardation time (10 s [45]).

It is worth noting that the extracellular space (ECS) consists of not only fluids but also proteoglycans, hyaluronan, and other extracellular matrix (ECM) components. These components can absorb and release water molecules quickly [3], and as the ECM is significantly softer than axons [92], explicit modelling of ECM components was omitted in this study.

A two-way coupling scheme was implemented to model fluid-structure interaction (FSI) between the drug fluid and axons. This iterative scheme updates axon deformation and drug flow status at each time step, accounting for large deformations and improving accuracy:

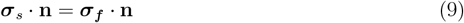

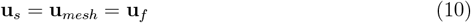

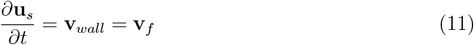

where **n** is the normal direction, **u** is the displacement, **v** is the velocity. The subscripts *s, mesh, f* denote the solid phase, mesh, and fluid phase, respectively.

In both flow directions, axons may come into contact. To account for this, penalty-based multibody contact algorithms were employed, assuming no friction or adhesion. Periodic boundary conditions were imposed on the computational domain to minimise size and boundary effects. All simulations were performed using the COMSOL Multiphysics platform. The finite element models passed mesh sensitivity tests.

### Permeability tensor derivation

Parametric studies were conducted to quantify the effects of pressure and pressure drop on tissue permeability by varying inlet pressure and pressure drop values. In real situations, the local pressure and pressure gradient change simultaneously. To derive the corresponding pairs of pressure and pressure gradient, Darcy’s Law was employed:

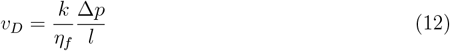

where *v*_*D*_ is the Darcy velocity, *κ* is the permeability, and Δ*p* is the pressure drop. The median permeability values measured from *ex vivo* infusion tests (*k*_∥_ = 1.5 × 10^−16^ m^2^, *k*_⊥_ = 0.7 × 10^−16^ m^2^) [64] were used in the calculations. The pressure range (0 ~ 3000 Pa) was selected based on tissue infusion experiments, with corresponding pressure gradients of 0 ~ 15 Pa in the parallel direction and 0 ~ 5 Pa in the perpendicular direction.

Fig. S3 illustrates the relationships between permeability and local pressure or pressure gradient. Parallel permeability (*k*_∥_) increases almost linearly with local pressure (Fig. S3a), while perpendicular permeability (*k*_⊥_) shows a parabolic increase (Fig. S3c), indicating greater sensitivity to higher local pressures. Although the pressure-parallel permeability relationship can be approximated by a linear function, slight non-linearities are present. Therefore, a parabolic function was used to describe the relationship between local pressure and permeability in both directions. The response of permeability to pressure gradient differs significantly between the two directions. Parallel permeability increases parabolically with the pressure drop (Fig. S3b), whereas perpendicular permeability exhibits exponential decay with the pressure drop (Fig. S3d). These relationships can be expressed as follows:

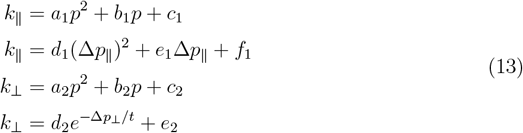

where *k*_∥_ and *k*_⊥_ are the parallel and perpendicular components of the local permeability tensor, respectively; *p* is the local pressure; Δ*p*_∥_ and Δ*p*_⊥_ are the local pressure drops in parallel and perpendicular directions, respectively; *a*_1_ ~ *f*_1_ are the coefficients of the parallel component, and *a*_2_ ~ *e*_2_, *t* are the coefficients of the perpendicular component.

By combining the *p* terms and Δ*p* terms in a first approximation, we can rewrite the permeability tensor as:

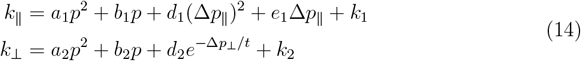

These equations were fitted to the *p*−Δ*p*−*k* data points obtained from the parametric sweep. As shown in Fig. S3e and S3f, the fitted models accurately describe the data, with coefficients of determination *R*^2^ exceeding 0.99 for both permeability components.

### *Ex vivo* sheep brain tissue infusion experiments

We briefly introduces the experimental details related to validate the tissue scale permeability tensor, while details can be found in Ref. [64]. Fresh ovine brains were collected from a local slaughterhouse. Samples were obtained from the CR region of the WM (see Fig. 2e), where axonal directionality is readily identifiable. The brains were sliced transversely and longitudinally to produce cylindrical samples oriented parallel and perpendicular to the nerve fibres. To facilitate testing, a plastic tube of the same dimensions as the cylindrical samples was used to wrap and support the tissues. A high-precision infusion-based experimental setup *iPerfusion* [93] was employed to infuse phosphate-buffered saline (PBS) into the samples. A stainless-steel needle (30G 1/2”; 0.3 13 mm) was inserted into the centre of the samples, as depicted in Fig. Fig. 2e. Pressure differences (ΔP), calibrated by a differential pressure transducer (Omega PX409) with an accuracy of 0.04 mmHg, were applied stepwise. Corresponding flow rates (Q) were recorded at steady state during each infusion step using a thermal flow sensor (Sensirion SLG150) with an accuracy of approximately 5 nL/min. Using Darcy’s Law and the measured pressure differences, flow rates, sample dimensions, and nerve fibre orientation, equivalent permeability values in the parallel and perpendicular directions were calculated, as shown in Fig. 2f. While these equivalent permeability values may differ from those obtained using traditional perfusion-based methods, the consistency of the measurement method ensures validity for the purposes of comparison with the present study. Compared to perfusion-based methods, which characterise homogenised permeability, the infusion-based method provides more localised permeability measurements [64]. Variations in experimental results arose due to the microstructural differences between tissue samples extracted from different ovine brains.

### Geometrical reconstruction of ovine brains from MRI

High-resolution and *in vivo* T1-weighted images of the ovine brains were acquired on a 1.5 Tesla Philips machine in San Raffaele Hospital in Milan with a voxel size of 0.667×0.667 mm and the slice thickness of 1.4 mm. The image datasets were then imported into the image processing software MIMICS (Materialise HQ, Leuven, Belgium), where the sheep brain and its ventricle can be segmented from the surrounding tissues and organs based on the signal intensity in three directions, as shown in Fig. 4a. By linking the segmentation on each image slice along the MR scan direction, the 3D geometry of the sheep brain and its ventricle were reconstructed. We then imported the geometry to COMSOL Multiphysics and generated the finite element mesh on the geometry. The mesh around the infusion site was refined to balance high accuracy and affordable computational burden.

### Anisotropy of ovine brains from DTI

Diffusion MRI (dMRI) enables the calculation of the diffusion coefficient of water molecules by applying a series of magnetic field gradients in different directions and measuring the corresponding random Brownian motion of water molecules within a voxel of tissue. The results reveal the brain’s microscopic architecture and connectivity by mapping the degree of restriction on water molecule movement within the tissue [94]. DTI, a specific type of diffusion-weighted MRI, provides information on both the directions and magnitudes of water molecule movement in each voxel as a 2nd-order diffusion tensor. This tensor comprises principal directions (eigenvectors) [*ē*_1_, *ē*_2_, *ē*_3_] and corresponding eigenvalues [*λ*_1_, *λ*_2_, *λ*_3_], as illustrated in Fig. 4b. By colouring voxels based on the direction of their greatest eigenvalues, the local orientation of nerve fibres can be visualised. For instance, in Fig. 4b, green indicates the greatest eigenvalue aligns with the dorsal-ventral direction, showing that the local orientation of nerve fibres in the green regions follows this direction. Similarly, blue represents the rostral-caudal direction, and red denotes the medial-lateral direction. Since water molecules diffuse faster along nerve fibres than transverse to them, DTI effectively indicates the direction of axonal bundles in the WM by connecting the directions of the greatest eigenvalues.

In this study, *in vivo* DTI datasets of the ovine brain were acquired with a voxel size of 22 mm and a slice thickness of 2 mm. The coordinates [x, y, z], eigenvectors [*e*_11_, *e*_21_, *e*_31_, *e*_12_, *e*_22_, *e*_32_, *e*_13_, *e*_23_, *e*_33_], and corresponding eigenvalues [*λ*_1_, *λ*_2_, *λ*_3_] in each voxel were combined and organised into a 15-row matrix. This matrix was imported into the whole-brain geometry to link local anisotropy to spatial coordinates, as shown in Fig. 4c. This dataset was used to define the local permeability tensor and diffusion tensor for whole-brain modelling.

### *In vivo* sheep brain infusion experiments

*In vivo* infusion experiments with four adult female sheep (6 hemispheres, 3 parallel infusion sets and 3 perpendicular infusion sets) were adopted to validate the accuracy of the newly developed framework. The experiments were carried on at Vita-Salute San Raffaele University, Italy, in the context of the EU’s Horizon EDEN2020 project. All animals were treated according to the European Communities Council directive (2010/63/EU), to the laws and regulations on animal welfare enclosed in D.L.G.S.26/2014. The experiment is briefly introduced here to show the rationale of the simulation set-up and the readers can refer to [95] for the details.

The sheep were anaesthetised and placed in the prone position on a 1.5T clinical scanner. DTI and MR images were taken to both guide the catheter implantation and brain geometry reconstruction. As shown in Fig. 3a, the ovine corticospinal tracts (CSTs, the white flocculent structure, also shown in Fig. 3f and 3f) were identified and visualised through the tractography technique. This helped to determine the entry point (E) and target point (T) for the catheter insertion, which enables the infusion can be both parallel and perpendicular to the local nerve fibres. A guide catheter with an outer diameter of 2.5 mm and an internal hollow was inserted from E to T via a tailored MRI-compatible ovine headframe and stereotactic system. After confirming that no haemorrhage or complication has induced by the surgery, the sheep were removed from the scanner and a specialised silica catheter with a diameter of 0.3 mm was then inserted through the guide catheter. For each animal, 10 *µ*L [69, 96] of a gadolinium-based solution (Prohance, 1:80 in saline, c = 0.5 mmol/mL) was infused with the rate of 3 *µ*L/min into the target positions by a syringe pump (Pump 11 Elite & Pico Plus, Harvard Apparatus, Holliston, Massachusetts, USA). Fig. 3b schematically shows the infusion setup.

After the infusion, the sheep were transferred to the scanner and 3DT1 scans were acquired at multiple time points (post-operation scans). The post-operation scans were co-registered to the 3DT1 baseline acquired right after the guide-catheter insertion. The aim was to extract the spatial distribution varying with the post-operation time of the infused Gd bolus, as shown in Figs. 3c and 3d. An in-house algorithm developed by the Vita-Salute San Raffaele team further enabled the extraction of the infused Gd as a 3D bolus and get their spatial relationship with the local nerve fibre, as shown in Fig. 3e. More details are given in the next section. The orientations of the Gd bolus and nerve fibres compared against the catheter are shown in Figs. 3f for the parallel infusion and 3g for the perpendicular infusion. The Gd boluses obtained at different time points were used to validate the whole brain modelling framework.

### Gd boluses reconstruction

To identify and map the areas of infused boluses in each time point (TP), we developed an automated method using a computer program called MATLAB. This method uses a “region-growing” algorithm, which starts from the brightest parts of the image and expands to include nearby areas with similar brightness, creating a mask that matches the edges of the bolus. We determined the brightest point (maximum intensity) in each bolus using a tool called ITK-SNAP, applying a small 3D mask that ensured the entire bolus was covered in the image. Another tool, FSL (https://fsl.fmrib.ox.ac.uk/fsl), was used to record this maximum intensity value. The darkest relevant point (minimum intensity) was identified by an experienced neuroradiologist, and these two values were used to calculate how much the brightness of the bolus fades over time (a measure called signal intensity decay). From this analysis, we found that on average, the brightness of the bolus fades by 33% (± 3.32%). Using this information, we programmed the algorithm to include only the areas that were at least 33% as bright as the brightest point of the bolus. The algorithm stopped growing the mask once the brightness dropped to 66% below the maximum intensity. Any areas dimmer than this were excluded from the bolus mask. This approach ensures that the mapped regions accurately represent the boluses observed in the images. Please refer to [95] for technical details.

### The multiscale modelling framework

According to the brain drug transport model built by Zhan et al. [97], the brain tissue can be divided into three compartments, namely the extracellular space (ECS), the neuron membrane (NM), and the neuron interior (NI). The concentration of free drugs that can act on target sites and cause a pharmacologically relevant effect (*C*_*F*_) and drugs that bind with proteins and are pharmacologically inactive (*C*_*B*_) is governed by the mass conservation equations: [97].

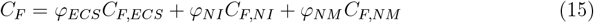

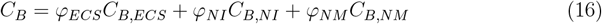

where *φ* refers to the volume fraction. It is assumed that there is no drug either being eliminated or associated with proteins on NM [98]. Free drug accumulation in the whole brain is determined by convective and diffusive transport in the IS, binding with proteins, cell uptake and elimination due to the loss to the blood circulatory system, physical degradation and metabolism. Therefore, the concentration of free drugs in the entire tissue is described by

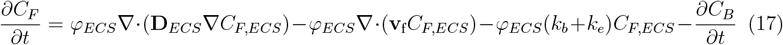

where **D** is the diffusion tensor of the drug particles; **v**_f_ is the velocity field of the drug; *k*_*b*_ and *k*_*e*_ are the elimination rate due to the blood drainage and degradation/metabolism, respectively. We further assumed that the free and bound drug concentration has a linear relationship [99], *i*.*e*.,

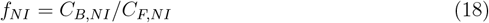

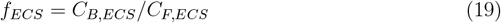

and the equilibrium of the free drug concentration is achieved among ECS, NM, and NI [100], *i*.*e*.,

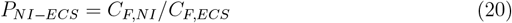

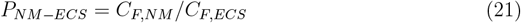

Therefore, Eq. 17 can be written as:

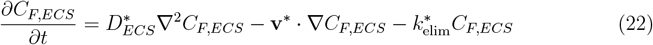

where 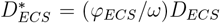 is the drug apparent diffusivity, **v**\* = (*φ*_*ECS*_*/ω*)**v**_f_ is the apparent interstitial flow velocity field of the drug fluid by assuming it as an incompressible fluid, and

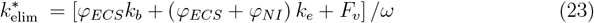

is the drug’s apparent elimination rate, and

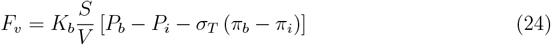

is the fluid flux from blood based on Starling’s law, and

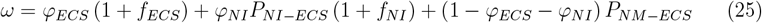

is a parameter that incorporates the drug abilities of local partitioning and binding, where *K*_*b*_ is the hydraulic conductivity of the blood vessel wall, 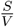 is the vascular density defined as the area of the blood vessel wall in the total tissue volume, *P*_*b*_ and *P*_*i*_ are the blood pressure and the interstitial pressure, respectively, and *π*_*b*_, *π*_*i*_ are the osmotic pressures of blood and interstitial fluid, respectively. *σ*_*T*_ is the averaged osmotic reflection coefficient for proteins in the blood.

In the *in vivo* experiments, the infused substance was Gd. Gd is stable and does not bind to protein *ex vivo*, it also has no detectable biotransformation or decomposition *in vivo* This means that the general mathematical framework can be significantly simplified as the Gd only exists in the ECS, *i*.*e. ω* = 1. In addition, the elimination rate due to the degradation/metabolism is zero, *i*.*e. k*_*e*_ = 0 in Eqs. 17 and 23.

The inlet boundary condition was assigned to the tip of the catheter. The inlet flow rate was 3.3 *µ*L/min with the initial pressure of 1853.18 Pa, as the average mean Intracranial Pressure (ICP) of sheep was measured as 13.9 ± 9.4 mmHg [101]. The inlet concentration was 0.5 mmol/mL. Outlet boundary condition (*P* = 1853.18 Pa, *c* = 0) was assigned to the outer boundaries of both the whole geometry and the ventricle. The other parameters that were applied in the simulations are summarised in Table S1. Given that the experimental time is much shorter compared to the tissue growth, drug transport properties and geometric parameters are assumed to be independent of time.

#### The information from the neuron and tissue scales

Since the drug delivery process is driven by convection and diffusion in CED, the apparent interstitial flow velocity field of the drug fluid (**v**\*) and the apparent diffusion coefficient of drugs in the brain ECS 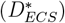 are two of the most important parameters in the above equation system.

As the brain WM was treated as a transversely isotropic fibrous porous medium, the apparent interstitial flow velocity field of the drug fluid can be obtained by applying Darcy’s Law:

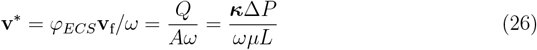

where *Q* is the flow rate, *A* is the cross-sectional area of the tissue, Δ*P* is the pressure drop over the tissue, *L* is the length of the tissue, the tissue refers to the region where the drug transport through, and

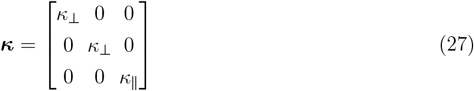

is the permeability tensor of the brain WM. See Eq. 14 and Fig. S3 for their formulas and values.

In terms of the apparent diffusion coefficient of drugs in the brain ECS 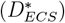, the particle dynamics model integrated with the microstructural geometry built in our previous work [40] has enabled us to characterise it. Since gadolinium-based solution was applied in the *in vivo* infusion experiments, the hydraulic diameter of gadobutrol (2nm) measured by Valnes et al. [102] was adopted, and the apparent diffusion coefficient was 2×10^−10^ m^2^/s.

However, the ***κ*** and 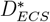 cannot be directly used in the whole brain model as they are the local parameters; their principle directions align to the orientation of the native nerve fibres and are defined in a local coordinate, as illustrated in Fig. 4c. To apply them in the whole brain, coordinate transformation is necessary. As the DTI data allows us to identify the eigenvectors [*e*_1_, *e*_2_, *e*_3_] and the corresponding eigenvalues [*λ*_1_, *λ*_2_, *λ*_3_] in each 2×2×2 mm^3^ voxel at the position with the global coordinates of [x, y, z], where

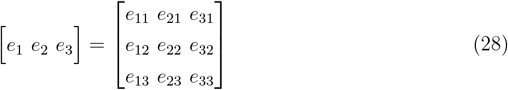

the apparent permeability tensor and diffusion coefficient tensor in the global coordinate can be obtained by:

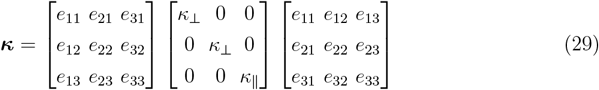

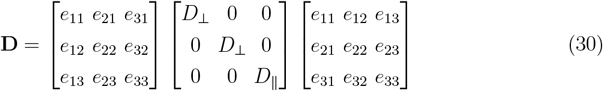

where the direction of *κ*_∥_ and *D*_∥_ align with the direction which has the largest eigenvalue.

All the simulations were conducted with the COMSOL Multiphysics platform and the finite element models have passed mesh sensitivity tests. For the sake of numerical stability, the rectangle function used to apply the inlet flow rate and concentration over the 3.3 min infusion time was smoothed.

## Acknowledgments

This project has received funding from the European Unions Horizon 2020 research and innovation program under Grant Agreement No. 688279. Daniele Dini would like to acknowledge the support received from the EPSRC under the Established Career Fellowship Grant No. EP/N025954/1, the Shell/RAEng Research Chair in Complex Engineering Interfaces (RCSRF2122-14-143), and the Medical Research Council (MR/Y008448/1).

Tian Yuan would like to acknowledge the support received from Great Britain-China Educational Trust award and the Medical Research Council (MR/Y008448/1).

