## Supplementary Information for "Decoding Brain Interstitial Transport In Vivo: A Fully Validated Bottom-Up Mechanistic Prediction Framework"

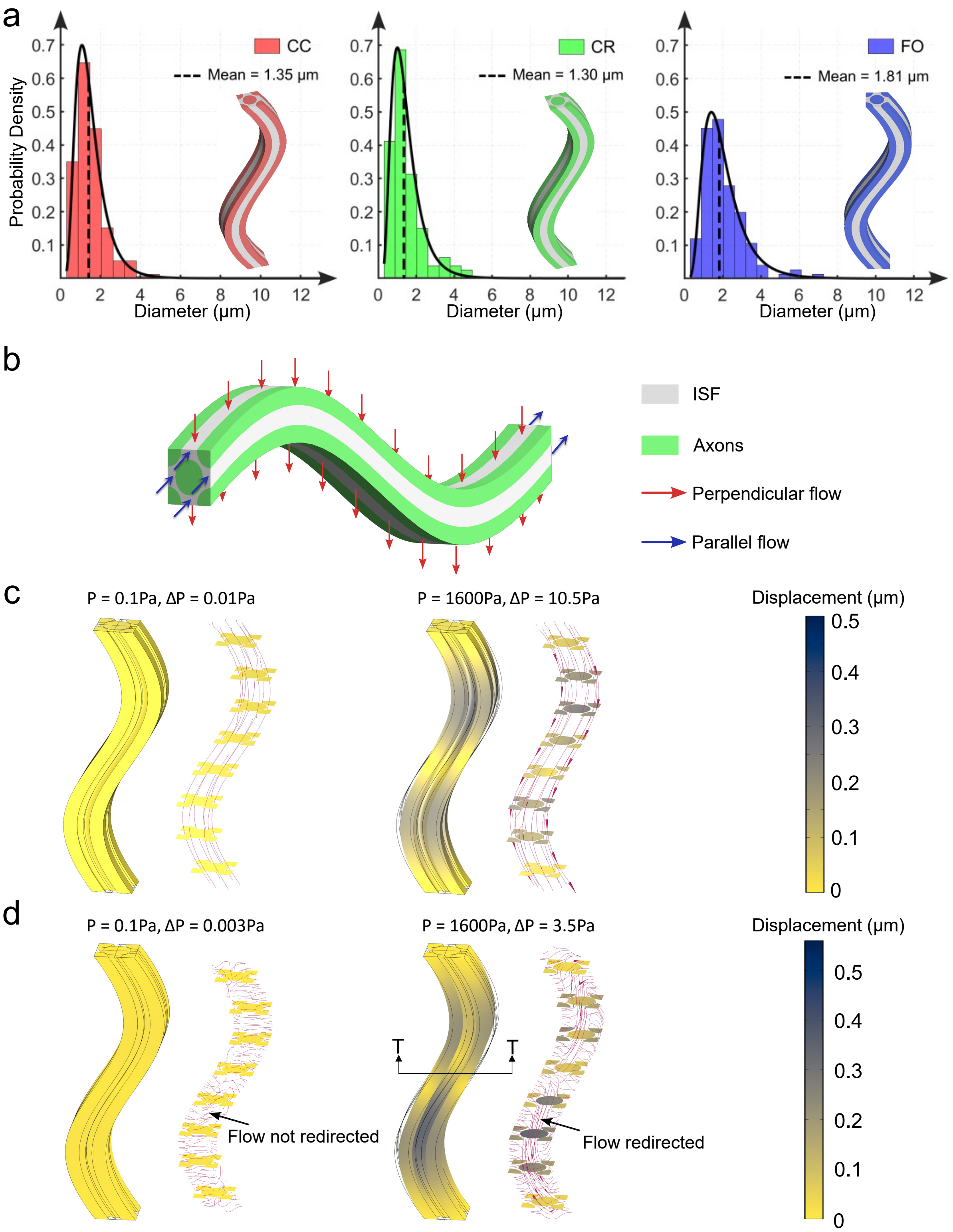


**Fig. S1**: **Modelling of the ISF-axons interactions**. **a**, The probability density of axon's diameters in CC, CR, and FO, respectively, obtained via FIB-SEM scanning [1]. The wavy structures are the 3D representative volume elements of the brain WM in the corresponding regions. **b**, 3D geometry and boundary conditions for the ISF-axons interaction modelling. **c**, ISF-axons interactions when fluid flows parallel to the axons. Simulations were run under different pressure and pressure drop pairs. In each group, the left-hand side panel shows the axons' deformation while the right-hand size panel demonstrates the flow status under the deformed flow pathway. In deformation figures, the wireframe shows the original(undeformed) shape of the axons. The legend on the rightmost side indicates the amount of deformation of the axons. In flow status figures, the arrows indicate the flow direction; their sizes are proportional to the flow velocity. **d**, ISF-axons interactions when fluid flows parallel to the axons. The figure descriptions are the same as that in **c**.

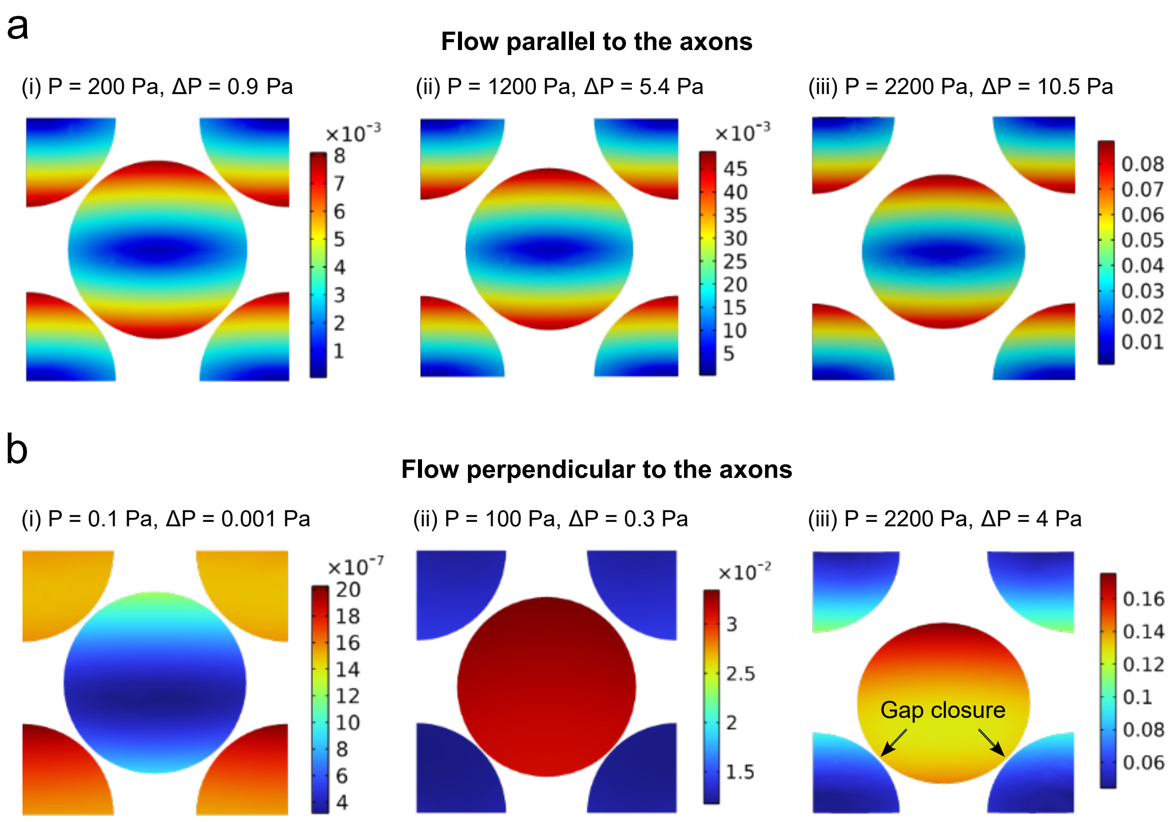


**Fig. S2**: **Axons deformation (unit: $\mu$m) in the T-T cross-section (see Fig. S1) caused by ISF flow in different directions.** **a**, Results of flow parallel to the axons. **b**, Results of flow perpendicular to the axons.

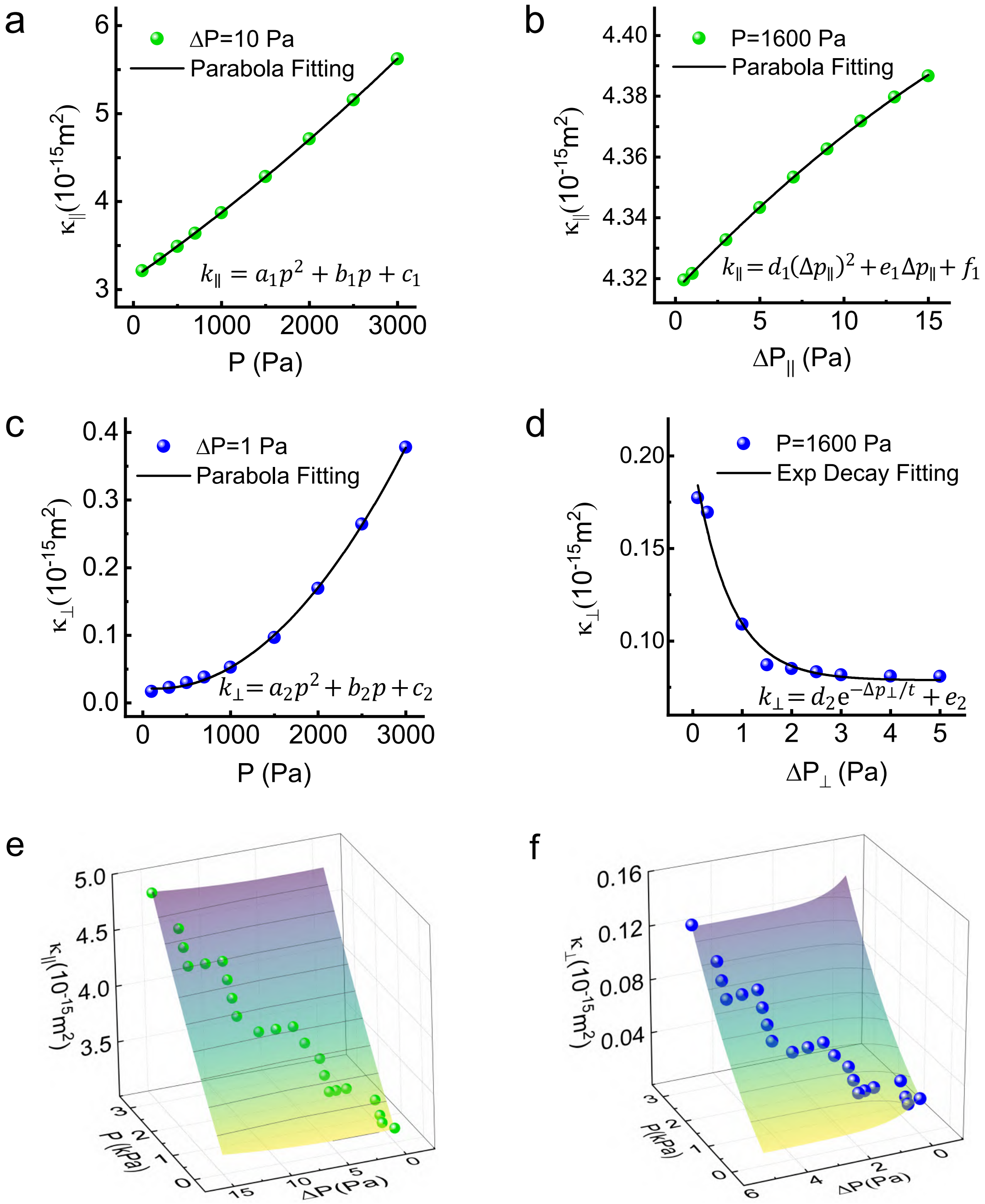


| $Eq.$ | $k_{\parallel} = a_1p^2 + b_1p + d_1(\Delta p_{\parallel})^2 + e_1\Delta p_{\parallel} + k_1$ |
|---|---|
| $a_1$ | $2.911 \times 10^{-23} \pm 4.875 \times 10^{-24}$ |
| $b_1$ | $4.849 \times 10^{-19} \pm 1.709 \times 10^{-20}$ |
| $d_1$ | $4.577 \times 10^{-19} \pm 1.839 \times 10^{-20}$ |
| $e_1$ | $-3.129 \times 10^{-15} \pm 3.255 \times 10^{-18}$ |
| $k_1$ | $3.129 \times 10^{-15} \pm 3.255 \times 10^{-18}$ |
| $R^2$ | 0.9999 |

| $Eq.$ | $k_{\perp} = a_2p^2 + b_2p + d_2e^{-\Delta p_{\perp}/t} + k_2$ |
|---|---|
| $a_2$ | $5.823 \times 10^{-24} \pm 6.236 \times 10^{-25}$ |
| $b_2$ | $2.059 \times 10^{-20} \pm 2.284 \times 10^{-21}$ |
| $d_2$ | $1.766 \times 10^{-17} \pm 2.410 \times 10^{-18}$ |
| $t$ | 0.713 ±0.021 |
| $k_2$ | $9.454 \times 10^{-18} \pm 1.984 \times 10^{-19}$ |
| $R^2$ | 0.99858 |

**Fig. S3**: **Derivation of permeability tensor. a**, Relationship between local pressure and permeability in the parallel direction. **b**, Relationship between local pressure drop and permeability in the parallel direction. **c**, Relationship between local pressure and permeability in the perpendicular direction. **d**, Relationship between local pressure drop and permeability in the perpendicular direction. **e**, Relationship between local pressure & pressure drop and permeability in the parallel direction. **f**, Relationship between local pressure & pressure drop and permeability in the perpendicular direction.

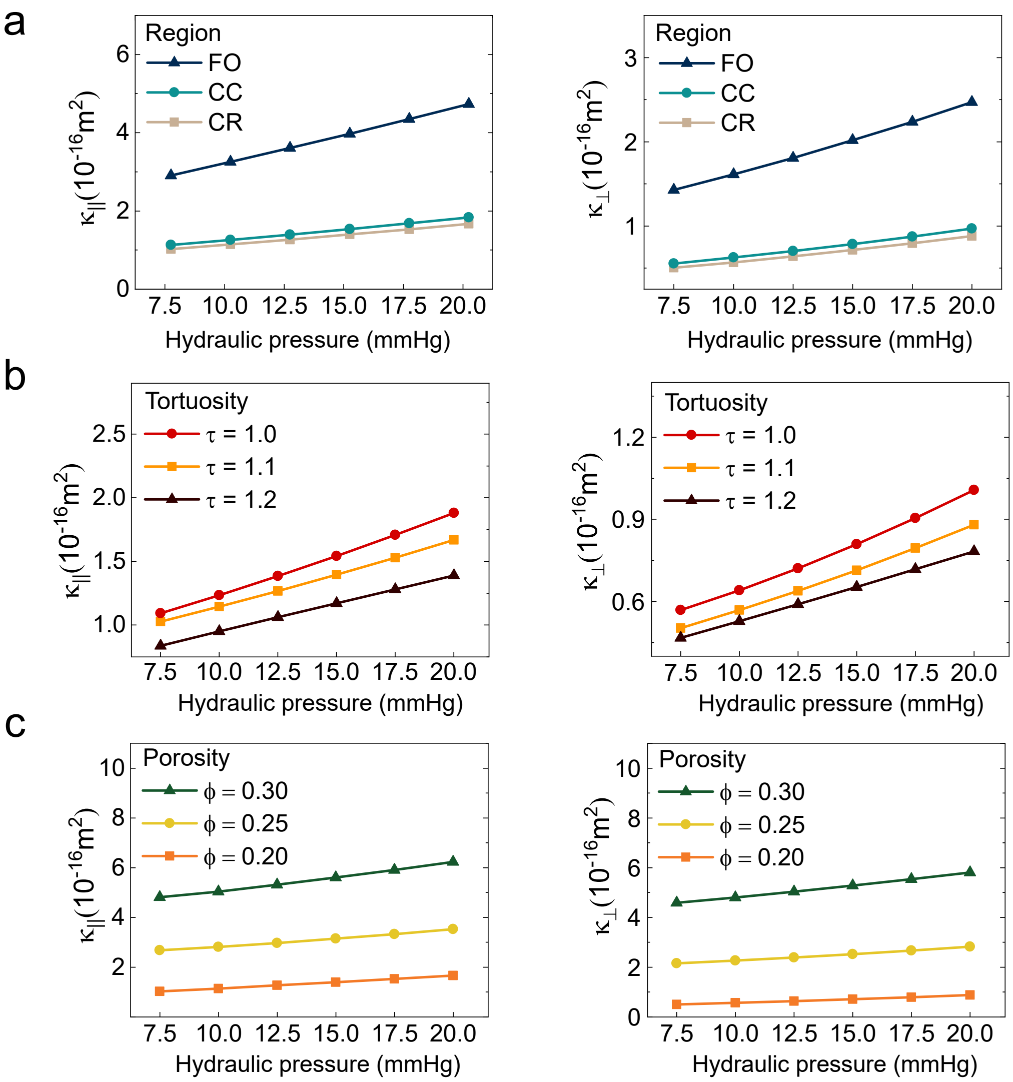


**Fig. S4**: **Variation of the pressure-dependent permeability tensors of WM. a**, Permeability tensors of FO, CC, CR under different hydraulic pressure. It indicates that transport efficiencies vary significantly in different regions. **b**, Tortuosity dependence of WM's permeability tensor. Axons in different regions may also have different tortuosity, but results imply that tortuosity does not dominate the change in transport efficiency. **c**, Porosity dependence of WM's permeability tensor. The normal average porosity of brain tissue is 0.2, but it can be enlarged by 50% to 0.3 when falling asleep. Results show that under this condition, permeability can be enhanced by 5 times under normal ICP (7.5 $\sim$ 15 mmHg), which means that the ISF bulk flow would be significantly facilitated thus helping to clear waste during sleep.

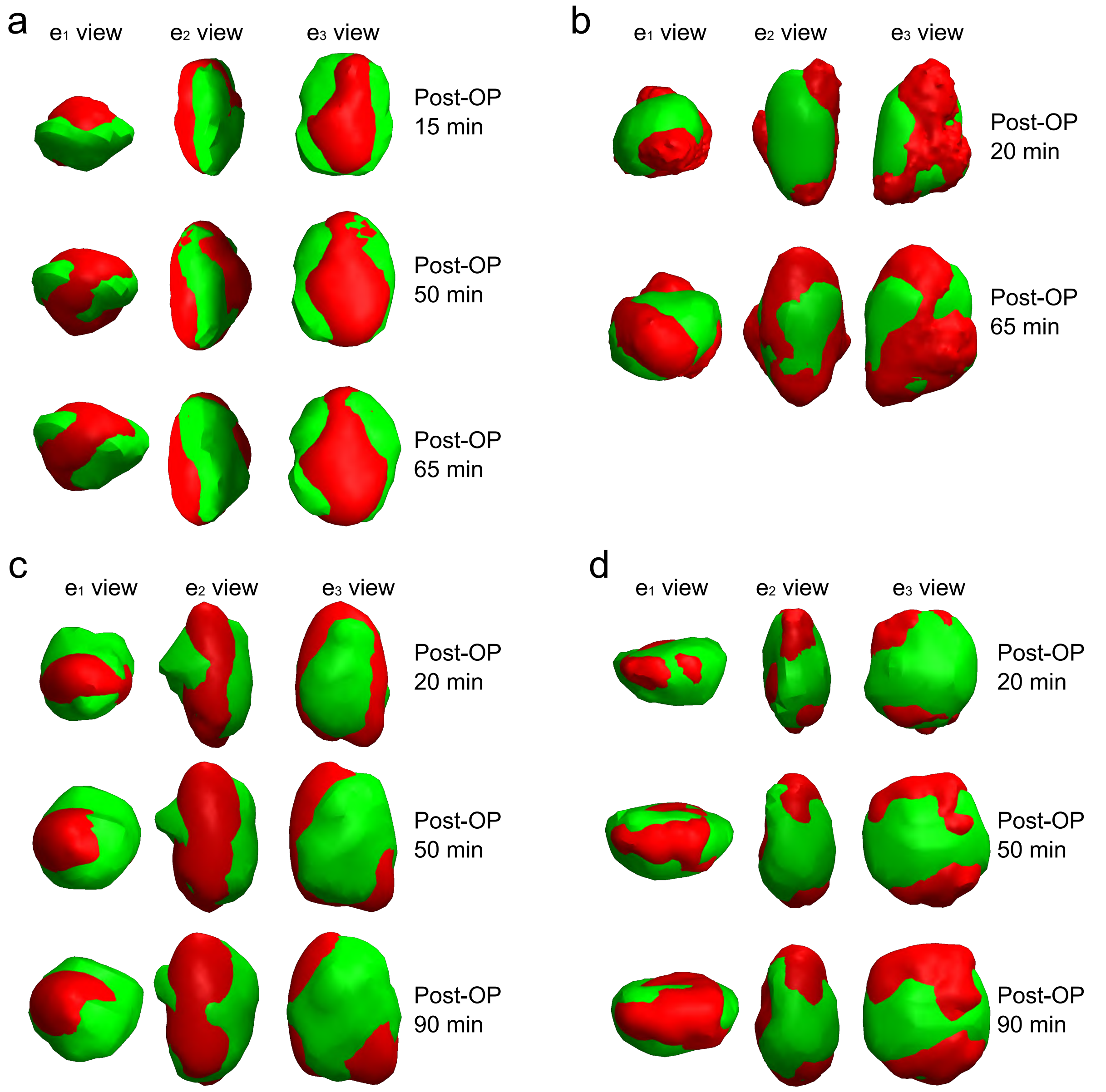


**Fig. S5**: **Comparison of the Gd bolus shape obtained by experiment (green) and simulation (red) in different groups and at different time points. a**, Group 2, parallel infusion. **b**, Group 3, parallel infusion. **c**, Group 5, perpendicular infusion. **d**, Group 6, perpendicular infusion.

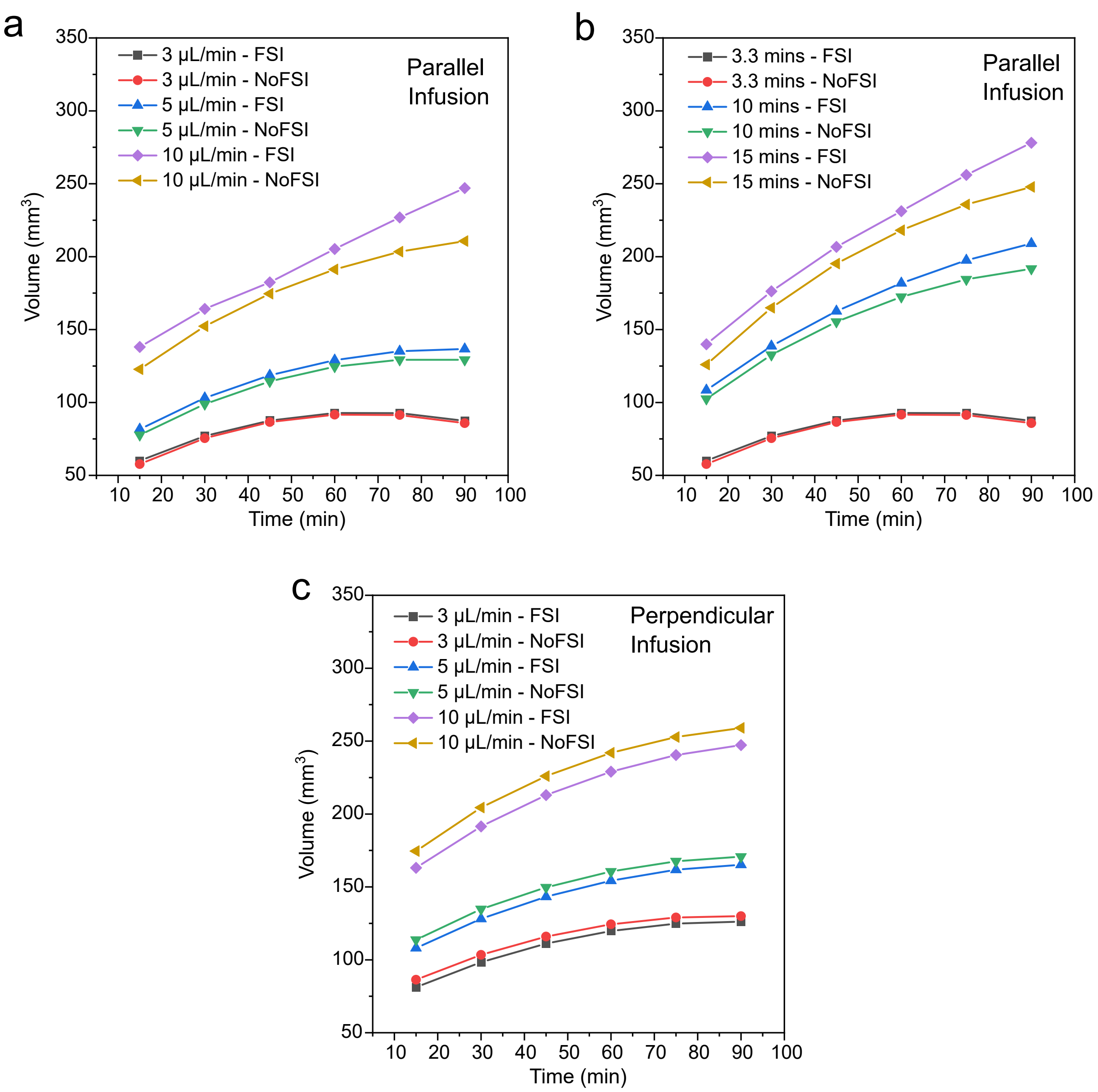


**Fig. S6**: **Effects of FSI on the Gd distribution in the brain after infusion.** The infusion duration was 3.3 minutes in both panels. **a**, The volume-time relationships under different infusion rates in the parallel infusion scenario. **b**, The volume-time relationships under different infusion durations in the parallel infusion scenario. **c**, The volume-time relationships under different infusion rates in the perpendicular infusion scenario.

**Table S1**: Parameters and values used in the whole brain modelling framework.

| Parameters | Descriptions | Values | Units | Refs. |
|---|---|---|---|---|
| $\varphi_{ECS}$ | Volume fraction of ECS | 0.2 | - | [2] |
| $\varphi_{NI}$ | Volume fraction of nerve interior | 0.65 | - | [3] |
| $\rho$ | Density of the Gd solution | 1000 | $kg/m^3$ | [4] |
| $k^*_{elim}$ | Apparent elimination rate of the Gd | 0.00276 | $min^{-1}$ | [5] |
| $K_b$ | Hydraulic conductivity of the blood vessel wall | $1.4\times10^{-13}$ | m/Pa/s | [3] |
| $P_b$ | Pressure in intravascular space | $4.6\times10^3$ | Pa | [3] |
| $S/V$ | Area of blood vessel surface per tissue volume | $7.0\times10^3$ | $m^{-1}$ | [3] |
| $\pi_b$ | Osmotic pressure of blood | $3.4\times10^3$ | Pa | [3] |
| $\pi_i$ | Osmotic pressure of interstitial fluid | $7.4\times10^2$ | Pa | [3] |
| $\sigma_T$ | Osmotic reflection coefficient of tissue | 0.91 | - | [3] |
